# Characterization and Expression Analysis of *Nitrate Reductase 6-1ABD* Gene in Hexaploid Bread Wheat Under Different Nitrogen Regime

**DOI:** 10.1101/2023.07.10.548320

**Authors:** Gayatri, Megavath Ravi, Harsh Chauhan, Ekta Mulani, Sachin Phogat, Karnam Venkatesh, Pranab Kumar Mandal

## Abstract

Nitrate reductase (NR) is the key rate-limiting enzyme of the nitrogen (N) assimilation process in plants, which has not been characterized in bread wheat under nitrogen stress, especially with respect to their homeologues. Total 9 *NR*s were identified and classified into 3 groups, which showed a close relationship with different wheat ancestors. The occurrence of N-responsive *cis*-acting regulatory elements like MYB, MYC, G-Box and GATA-motif confirmed their N-responsiveness. Expression of all the three groups of *NR* under N-stress revealed *NR 6-1ABD* group to be the most N-responsive, which was characterized further in detail. The study was carried out in two genotypes contrasting for their N-responsiveness (HD 2967: Highly responsive to applied N, and Choti Lerma: Less responsive to applied N) selected on the basis of field evaluation. Homeologous differences within a genotype were found much more than the genotypic differences of a specific homeologue coding sequence. Among the three homeologues, though *NR 6-1D* homeologue was found most responsive to N-stress, the contribution was maximum for this homeologue followed by *NR 6-1A* and least by *NR 6-1B.* We found that the expression of homeologues was linked to the presence of N-responsive *cis-* elements. All the homeologues of *NR 6-1* in Choti Lerma were found less responsive to N-stress, in comparison to HD 2967, which might also be linked to N-use efficiency. Homeologous expression of *NR 6-1ABD* revealed negligible contribution of *B*-homeologue to N-stress. Homeologous differences of *NR 6-1ABD* was found much more than the genotypic differences. Hence, our study on wheat *NR* will be helpful in manipulating the specific homeologue of the *NR* gene in the future.

## Introduction

Wheat (*Triticum aestivum* L.) is the primary source of calories (>45%) and nutrition for the human population as well as essential for animal feed (Alipour et al. 2017; Hawkesford and Griffiths 2019; Mălinaş et al. 2022; Rawal et al. 2022). Consequently, excess use of nitrogen (N) fertilizer is applied to meet the higher demand for wheat production (Hirel et al. 2007). However, the N use efficiency of crop plants is low (around 33%) and more than 50% of the expensive applied N has been lost to the soil system (Gaju et al. 2011), leading to negative repercussions on the environment and human health (Foulkes et al. 2009; Mălinaş et al. 2022). Therefore, it is requisite to identify and develop novel strategies to enhance the efficient use of N to reduce the over-dependence on N fertilizers (Hawkesford 2014). Nitrogen use efficiency (NUE) consists of two components; nitrogen uptake (NUpE) and the other is nitrogen utilization efficiency (NUtE) (Moll et al. 1982; Mălinaş et al. 2022). N assimilation, transport, and remobilization are the three major components of NUtE (Li et al. 2017), which determine the yield. Hence, for the concept ‘low nitrogenous fertilizer but high yield’, NUE enhancement of wheat has now become a major challenge for wheat improvement programs. NUE is further studied as agronomic NUE and biological NUE (Sharma et al. 2021). Inherent plant NUE needs to be addressed biologically, and to date reports on improved NUE in wheat are very few, probably because of the very complex genome and its regulation. Also, the molecular basis of NUE is not yet fully understood and the biotechnological approaches have met with limited success so far (Sharma et al. 2021).

The first step of nitrogen metabolism in plants is the uptake of the inorganic form of nitrogen from the external environment into the cell. The major form of N taken up and assimilated by most of the crops under normal field conditions is Nitrate (NO_3_^-^) (Rosales et al. 2011; Wang et al. 2011; Garcia-Oliveira et al. 2013; Zhao et al. 2013). Hence, the availability of nitrate determines the crop growth, development, and grain yield in wheat (Kumari 2011; Nikolic et al. 2012). The plasma membranes of root epidermal as well as cortical cells have nitrate transporters, which are involved in the acquisition of nitrate molecules from the soil in an active process (Fan et al. 2017). Cytosolic nitrate reductase (NR) first reduces the nitrate up-taken by the plant and then chloroplastic nitrite reductase (NiR) converts the nitrate to ammonium (Stitt et al. 2002; Rolly and Yun 2021). This ammonia is assimilated as glutamate, the first amino acid, by the two enzymes glutamine synthetase (GS) and glutamate synthase (GOGAT), which work in a cyclic manner (Lea and Miflin 2003; Good et al. 2004; Gayatri et al. 2021). Thus, NR is considered a key and rate-limiting enzyme in the complex process of nitrate assimilation and utilization (Campbell 2001; Tang et al. 2022). NR also influences nitrate transporter proteins like NRT1.1 and NRT2.1, in addition to its key role in N assimilation (Lejay et al. 1999). A few reports of some higher plants also described a significant positive correlation of the parameters like protein content, growth, yield, and nitrogen status with NR enzyme in seeds and leaves (Hageman 1979; Srivastava 1980; Guerrero et al. 1981). Using biotechnological approaches like single gene transgenics, there have been several attempts to improve N metabolism in plants but with little improvement of NUE (Raghuram and Sharma 2019; Sinha et al. 2020; Sharma et al. 2021). However, there is broad scope to work with the genes involved in N metabolism. Allohexaploid bread wheat (*T. aestivum*) has three genomes A, B, and D with 2n = 6x = 42 (Marcussen et al. 2014; Lv et al. 2021). Hence, generally, three copies of a gene (homoeologous genes) are expected in bread wheat (Shitsukawa et al. 2007). The expression of numerous genes at their homeologous level has been reported in wheat (Kashkush et al. 2002; Himi and Noda 2004; Bottley et al. 2006; Shitsukawa et al. 2007; Hu et al. 2011; Gayatri et al. 2019, 2021). However, environmental conditions, the growth stage of the plant, and also the tissue determine the homoeologue-specific expression of any gene in bread wheat (Mochida et al. 2004; Bottley et al. 2006; Gayatri et al. 2019, 2021).

NR is the first enzyme which starts converting nitrate in to ammonia in the process of N assimilation and has been studied in detail in many crops. However, in wheat most of the reports are about the NR enzyme activity (Minotti and Jackson 1970; Blackwood and Hallam 1979; Sherrard and Dalling 1979; Botella et al. 1993; Carillo et al. 2005) and a few reports are about the *NR* gene expression (Chandna et al. 2012; Balotf et al. 2016). Bread wheat being a hexaploid crop, harbours three genome species in it. Therefore, most of the genes were assumed to be present in three copies which are called as homeologues. Many of these genes in bread wheat showed homeologue biased expression (Himi and Noda 2004; Nomura et al. 2005; Shitsukawa et al. 2007). So far, there is no information related to homeologue specific expression of *NR* gene along with their genomic information under different levels of nitrogen treatments. Hence, to study the homeologous expression of *NR* gene in wheat, first requisite was to find out the copies and homeologous set among the different copies of *NR*. Genome-wide identification of *NR* gene revealed nine copies are present in hexaploid wheat (Hurali et al. 2022). After confirming the same, we carried out (1) expression profiling of homeologous set (categorized on the basis of similarity) to check their N-responsiveness, and (2) homeologue-specific *NR* coding sequence cloning and analysis of the most responsive set of the genes, followed by their expression and contribution in two wheat genotypes that are contrasting for their N-responsiveness. Overall our objective and focus of this study was to reveal the homeologue specific hidden information of *NR* genes in bread wheat and their expression pattern as well as contribution at different nitrogen regime, which is so far not available. This study also would provide the foundation for functional characterization of each homeologue of *NR* in future to improve the N-assimilation process.

## Methods

### Identification and characterization of the *NR* genes using Chinese Spring Wheat Genome

To identify the *NR* genes in the wheat genome, we used our previously submitted wheat *NR* sequence (GenBank: KY244026.1) and used *EnsemblPlants* and IWGSC databases for BLASTN analysis as mentioned in Gayatri et al. 2022 (Gayatri et al. 2021). Thereafter, genomic, CDS, and amino acid sequences for all the *NR* genes were downloaded from these databases. Conserved domains and other information (like physio-chemical properties and subcellular localization) related to NR proteins were predicted using the Pfam database, ExPASy’s ProtParam tool, and CELLO2GO as mentioned in Gayatri et al. 2022 (Gayatri et al. 2021).

Wheat-URGI BLAST (https://wheat-urgi.versailles.inra.fr/Seq-Repository/BLAST) was used to predict the chromosomal position of each *NR* gene onto the wheat genome. The *NR* genes were named sequentially based on their position on the chromosomes, starting from *NR.* DNAMAN software was used to make the homeologous group of the most similar *NR* genes.

### *Cis-*acting elements in the *NR* promoter regions

For the promoter region, the upstream 1000 bp of each *NR* gene sequence was extracted from the *EnsemblPlants* database. Then this upstream region was submitted to the PlantCARE software(Lescot et al. 2002) to identify the *cis*-elements in the putative promoter regions.

### Phylogenetic Analysis and Homology with *NR* from other plants

The orthologous *NR* protein sequences (File S1) from diverse monocot and dicot plant species were obtained based on the sequence identity >65% using the NCBI database and an unrooted phylogenetic tree was constructed with 1000 replicates bootstrap parameters using the neighbor-joining method using CLC Genomics workstation 12. *NR* genes of wheat were compared with *NR* genes of other plant species and their genomic positions were recorded. The Galaxy Circos tool was used to identify the relationship between species and genomic positions in a circular form.

### Plant material, growth in the field and measurement of NUE related parameters

Several diverse wheat genotypes including HD 2967 and Chotio Lerma were grown during the Year 2013-14 with two doses of applied nitrogen (150 kg/ha: denoted as Y1-150 as and 0 kg/ha: denoted as Y1-0) under standard package of practices (Bharati et al. 2022) at ICAR-Indian Institute of Wheat & Barley Research (ICAR-IIWBR), Karnal, India.

During the year 2014–2015, we have grown these genotypes including HD2967 and Choti Lerma with four different levels of applied nitrogen i.e., 150 kg/ha (denoted as Y2-150); 120 kg/ha (denoted as Y2-120); 90 kg/ha (denoted as Y2-90) and 0 kg/ha (denoted as Y2-0) with standard package of practices. Both the Year the average available nitrogen in the field was 150 Kg/ha. The whole experiment was conducted in three replications.

### Selection of contrasting genotypes

At maturity, grain yield (GY) was determined from 1-m^2^ sampling area of each subplot. The N content of the dry biomass was determined using the Kjeldahl method. The GY and total N content of dry biomass were measured and used for calculating other derived parameters i.e Grain N Y/P, N at Harvesting, N-uptake, NUE (Bharati et al. 2022). Finally, values of NUE i.e [(GY/P)/((Soil N+Applied N)/P) were considered for the selection of most contrasting genotypes.

### Growth of plants under control conditions

Seeds of both genotypes were surface-sterilized using 0.5% (w/v) sodium hypochlorite solution and then used for germination on water-soaked germination paper at 25 ± 2 °C for 3 days in dark conditions. The hydroponic system (500ml plastic pot) containing 1X MS (Murashige and Skoog) solution without N was used as described by Gayatri et al 2022 (Gayatri et al. 2021) for the growth of wheat seedlings. N was added in the form of calcium nitrate (Ca (NO_3_)_2_.4H_2_O) in MS solution at two concentrations (8 mmol/L for N-optimum and 0.08 mmol/L for N-stress) (Gayatri et al. 2019, 2021). All the experiments were carried out in a controlled environment at 150–200 μmol photon/m^2^/s light intensity, 70% relative humidity, 8/16 dark/light hours, and 25 ± 2°C for 21 days and with three replications of each pot (Five seedlings per pot). Chronic and transient, two kinds of nitrogen stress were imposed for the present study. For chronic stress, seedlings were grown under N-stress conditions for 21 days. On day 21, the shoot and root tissues of the seedlings were harvested for measurement of biomass, chlorophyll and carotenoids content. For transient stress, seedlings were initially grown under optimum N condition for 14 days and then transferred to N-stress condition. Shoot tissues of the seedlings were harvested on 1 day (15 days After Transfer (DAT)), 4 days (17 DAT), and 7 (21 DAT) days after the imposition of N-stress. For further molecular studies, shoot tissues were harvested, pooled, and stored at −80°C. Each biological replicate consisted of pool of five plant samples, which were grown in one pot. Three such biological replicates were used for each treatment.

### Measurement of biomass parameters and chlorophyll pigments

Different biomass parameters i.e. shoot length (SL), root length (RL), shoot fresh weight (SFW), root fresh weight (RFW), root dry weight (RDW) were measured according to the standard protocol (Sinha et al. 2015, 2018). Chlorophyll (Chla, Chlb, TChl) and carotenoid contents were also estimated using fresh shoot tissues (Hiscox and Israelstam 1979) of both genotypes.

### Determination of *in vitro* NR activity

To determine the NR activity, total soluble protein was measured using the method described by Lowry et al., 1994; from 21 days old stored shoot tissues of both genotypes. Further, stored shoot tissues were powdered with a mortar and pestle and solubilized in phosphate buffered saline pH 7.4. The extract was centrifuged at 6000 g for 15 min and the supernatant was stored at -70°C. NR activity was examined by the reduction of nitrate to nitrite (Nikolic et al. 2012; Fan et al. 2017). One unit of NR activity is defined as the production of 1μM nitrite per min. Specific activity is described as a unit of NR activity divided by gram of total protein.

### Measurement of N contents

Dried shoot and root samples were used to determine the N content of both genotypes using FOSS NIRS™, DS2500 analyzer (Denmark)

### Total RNA isolation and cDNA synthesis

Total RNA extraction using the RNeasy Plant Mini Kit (Qiagen, Hilden, Germany) and cDNA synthesis using SuperScript®III First-strand Synthesis SuperMix (Invitrogen™, USA) kit were done from stored shoot tissues DNA as mentioned in Gayatri et al. 2019 and 2022 (Gayatri et al. 2019, 2021).

### Quantitative real-time PCR (qPCR)

*NR 6-ABD-1*, *NR 6-ABD-2*, *NR 7-AD-4A* sequences were used for designing the three sets of qPCR primers (Table S1b) based on their common sequences. Further, homeologue-specific primers (Table S1c) for qPCR were designed based on the nucleotide polymorphisms found in the cDNA sequences of the three homeologues for *NR 6-ABD-1*gene (Table S1a) (Gayatri et al. 2019, 2021). Actin gene (KX533928.1) was used as endogenous control. qPCR was performed and relative fold changes in the expression of N-stress samples over control were calculated according to Gayatri et al. 2019 and 2022 (Gayatri et al. 2019, 2021)The specificity of each homeologue-specific primer pair was determined by sequencing the amplified products (Gayatri et al. 2019, 2021).

### Cloning of full-length cDNA sequences of wheat *NR 6-ABD-1* homeologues

Homeologue-specific PCR primers (Table S1) for the *NR 6-ABD-1*gene were designed using the method explained by Gayatri et al. 2022 (Gayatri et al. 2021). The full cDNA of each *NR* homeologue was amplified as a single fragment using sequence-specific primers. High fidelity *Taq* polymerase (PrimeSTAR GXL DNA polymerase, Takara) was used in the preparation of the PCR reaction mixture followed by the PCR program according to Gayatri et al. 2022 (Gayatri et al. 2021). The PCR products were then cloned into a pJET1.2 vector (CloneJET, Thermo Scientific™, USA) and three clones for each PCR product were sequenced by Sanger sequencing (Genotypic Technology Pvt Ltd).

### *In silico* analysis of cloned *NR 6-ABD-1* homeologous cDNA sequences and their putative proteins

ORFs of the cloned *NR* homeologues cDNA sequences were predicted using the Open Reading Frame (ORF) finder (http://www.ncbi.nlm.nih.gov/gorf/gorf.html) and then the EXPASY tool was used to predict their respective amino acid sequences. ExPASy proteomics server was used to predict the theoretical molecular weight (MW) and isoelectric point (pI) of all the putative NR proteins. Further, for comparison and multiple sequence alignment (MSA) of cloned cDNA sequences as well as their putative NR proteins, DNAMAN software was used. To predict the potential impact of amino acid changes on the final protein structure (Ng and Henikoff 2003) as well as on the protein function, SIFT software was used.

### Statistical Analysis

All experiments reported in this study were done with three biological three technical replicates. MSTATC was used to perform the statistical analysis of data obtained in the present study. Mean values were presented for all the data with an error bar (standard error of means). T-test was performed to show the NUE contrasting nature of both genotypes. A least significant difference (LSD) at 5% was calculated for the significance of the treatment effect and further, a range test was performed to show the level of significance between and among the treatments in each experiment.

## Results

### Genome-wide identification and nomenclature of *NR* gene sequences in wheat (Chinese spring) genome

We initially confirmed the presence of nine copies of *NR* gene from the Chinese Spring wheat genome (Hurali et al. 2022). The *NR* genes were grouped in different groups to identify their homeologues set (indicated in different colours in Fig.1) showed an uneven distribution across the A, B, and D sub-genomes. In first group TraesCS6A02G326200, TraesCS6B02G356800 and TraesCS6D02G306000; in the second group TraesCS6A02G017500, TraesCS6B02G024900 and TraesCS6D02G020700 and in third group TraesCS7A02G078500, TraesCS7D02G073700 and TraesCS4A02G376700 were clustered together.

**Fig. 1.**
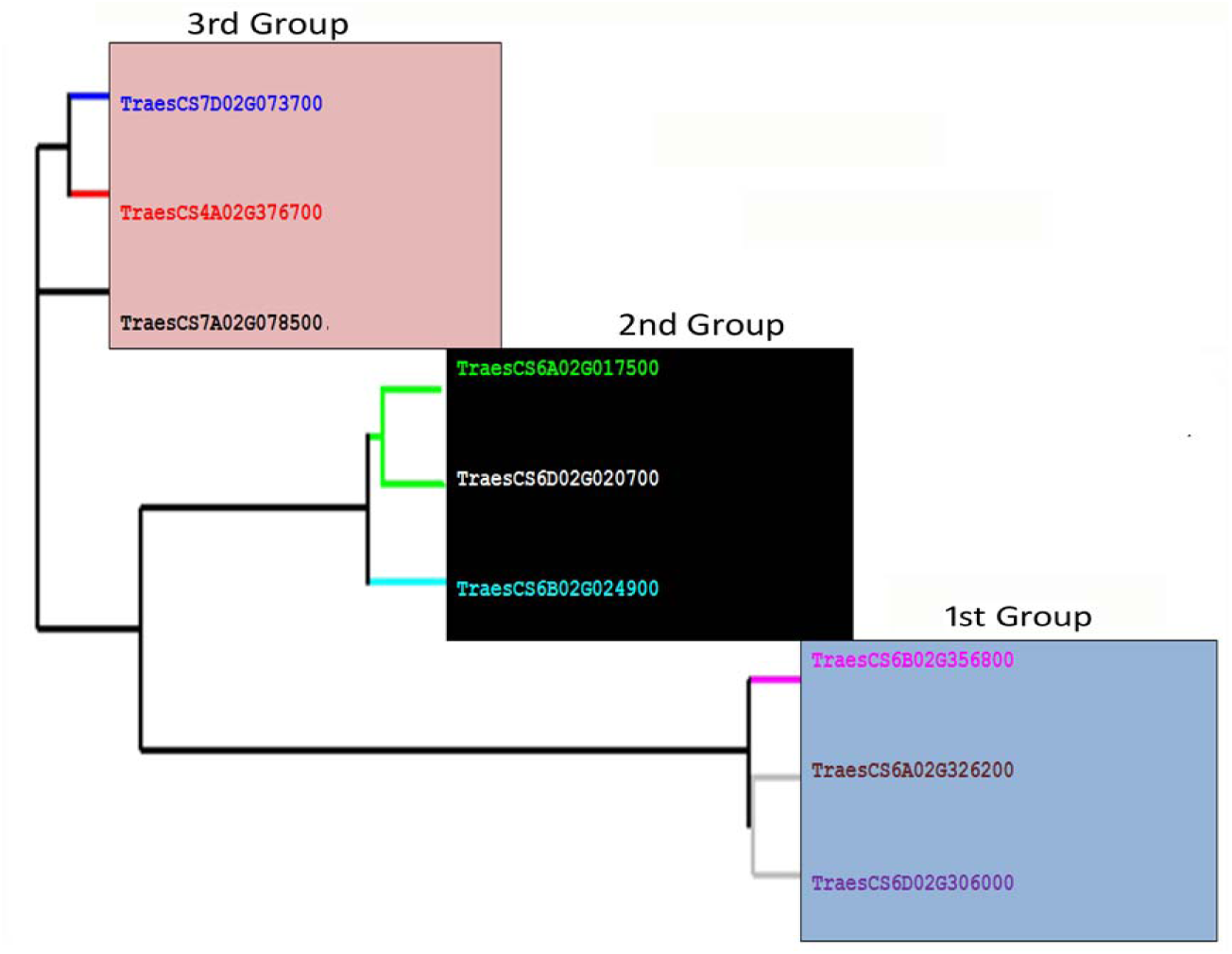
Classification of nine *NR* genes in to three groups on the basis of their similarity using DNAMAN The sequences were named *NR 6-1ABD* (first group, homeologues as *NR 6-1A*, *NR 6-1B*, *NR 6-1D*), *NR 6-2ABD* (second group, homeologues as *NR 6-2A*, *NR 6-2B*, *NR 6-2D*) and *NR 7AD-4A* (third group homeologues as *NR 7A*, *NR 7D*, *NR 4A*). This result also reveals that the B genome has only two copies of the genes in Chromosome 6, whereas D has three copies, two in chromosome 6 and one in chromosome 7 and A has four copies of the *NR* gene, two in chromosome 6, one each in chromosome 4 and chromosome 7.

### Phylogenetic and Collinearity Analysis between Wheat and other plant *NR* Genes

To further evaluate the phylogenetic relationships and collinearity analysis of *NR*s and *NR*s from other plants, we selected *NR* protein sequences different monocot and dicot species *NR* protein sequences having more than 65% similarity were used for the phylogenetic tree construction. As shown in Fig. 2, *NR*s from the selected plant species were divided into four subgroups. The largest subgroup is (IV) containing 6 *NR* sequences (Fig. 2). Wheat *NR* sequences were found to be most closely related to its progenitors’ sequences as well durum wheat along with *Hordeum vulgare* and *Sorghum bicolor*.

**Fig. 2.**
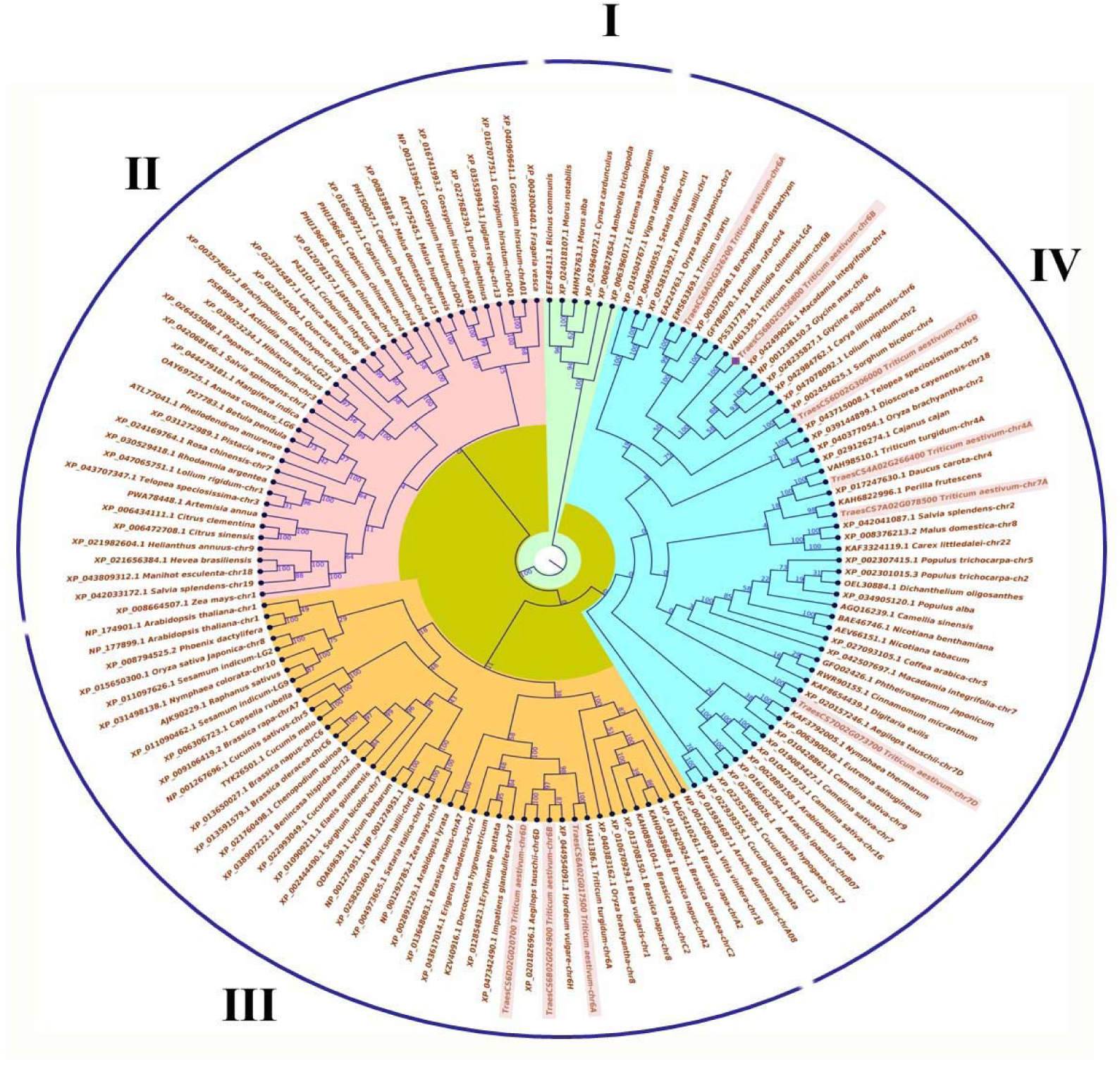
Phylogenetic relationship among *NR* genes of *T. aestivum* (highlighted) and other plant species. The unrooted phylogenetic tree was constructed using MEGA 6.0 Maximum-likelihood method with 1000 replicates bootstrap analysis. The tree is divided into IV subgroups

From phylogenetic analysis, we have found wheat *NR* close similarity with its progenitors and to investigate the change of *NR* gene number within a subgenome due to transition from diploidy to hexaploidy. For this, we have identified and extracted the *NR* gene from the diploid progenitors (*Triticum urartu* and *Aegilops tauschii*) and tetraploid wheat (*Triticum turgidum*) using *EnsemblPlants*.

To gain a more detailed outlook on the possible information on evolutionary divergence of the *NR* genes, the collinearity analysis was performed using the Circos tool among *T. aestivum*, *T. urartu, A. tauschii,* and *T. turgidum/durum*. From collinearity analysis, we illustrated that *NR* orthologs of *A. tauschii* were located on 6D and 7D chromosomes (two on 6D, and one on 7D) (Fig. 3). Furthermore, two *T. urartu* orthologous *NR* genes were found located on the 7A chromosome and ungrouped scaffold. In *T. turgidum,* 6 orthologous *NR* genes (two on 6A and 6B, one on 4A, and one on 7A) were found. On the whole, intact collinearity among the *NR* genes of bread wheat and its relatives suggested the conservative evolution of this gene during the formation of hexaploid wheat.

**Fig. 3.**
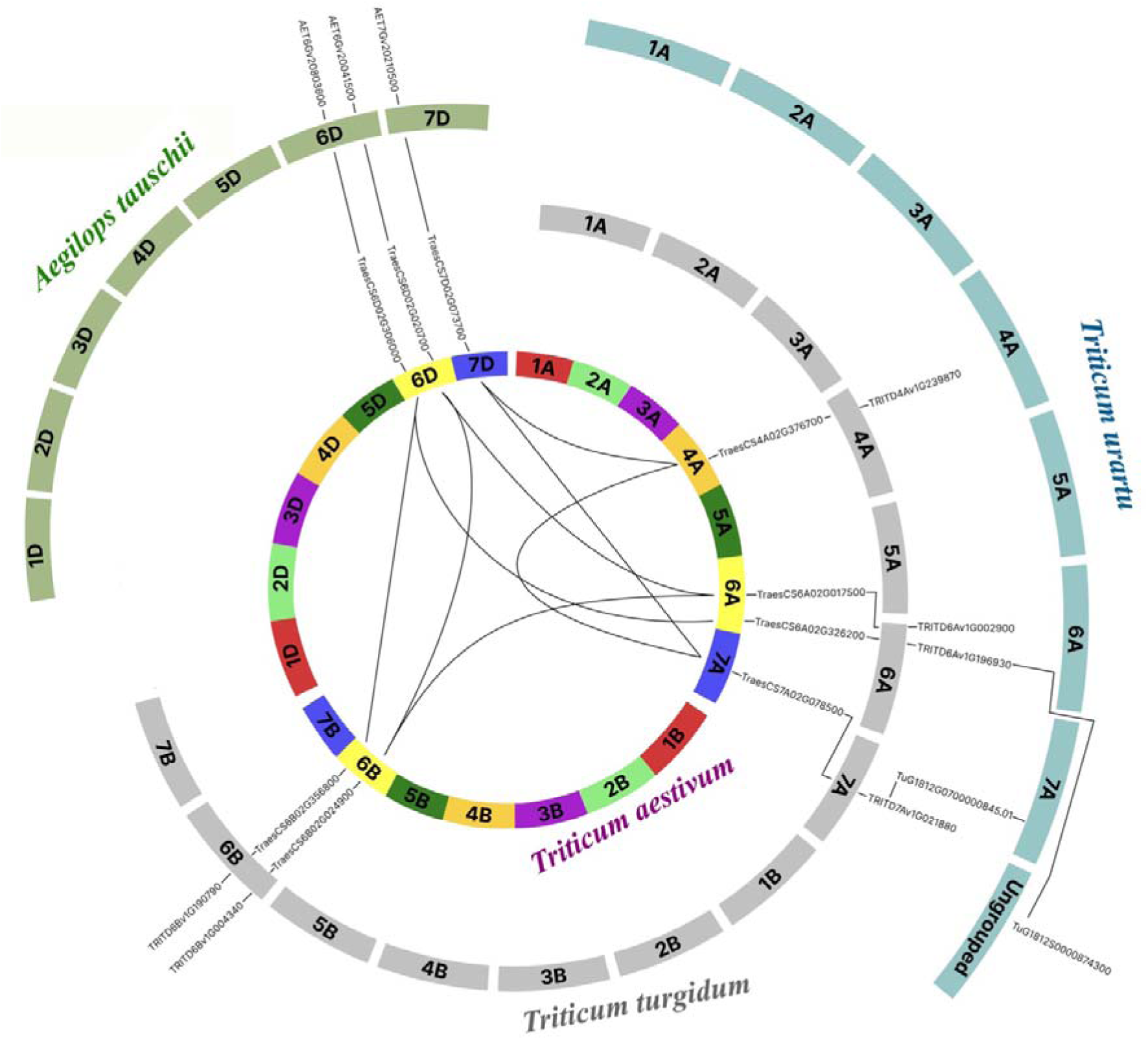
Collinearity Analysis of *NR* genes. Muticolour, sea-green, light green, and grey-coloured blocks represent the chromosomes of *T. aestivum*, *T. urartu, A. tauschii,* and *T. turgidum* respectively. The black color lines indicate orthologous as well as homeologous *NR* genes on different chromosomes. GeneIDs starting with *Trae, Tu, AET, TRITD* represent genes for *T. aestivum*, *T. urartu, A. tauschii* and *T. turgidum*

### Prediction and analysis of *cis-*acting elements in the promoter regions of *NR genes*

*Cis*-acting elements predicted in the promoter regions of the *NR* genes are associated with environmental stress, hormone response, light response, development, promoter and enhancer elements, site-binding elements, and others (Fig. 4b). To understand the potential mechanism via which *NRs* respond to biotic and abiotic stresses, we used PlantCARE to analyze the *cis*-regulatory elements in the promoters. Transcription-related *cis*-elements–including the TATA-box and CAAT-box were found in all the *NR* genes with maximum abundance (Fig. 4a). Our data revealed that almost all *NR* promoter regions contained at least one plant hormone-response element, including gibberellin-response element GARE-motif, ethylene-response element ERE, salicylic acid-response element TCA, jasmonic acid (MeJA)-response element CGTCA-and TGACG-motif and abscisic acid-response element ABRE. Light responsive elements such as ACE, G-Box, Box 4, TCT-motif, MRE, Sp1, AE-box, etc. were also identified. The promoter region also contained stress response elements, such as low-temperature stress-response element LTR, drought-response element MBS and low N stress response elements MYB, MYC, G-box, GATA-motif (Lea et al. 2007; Zhang et al. 2015; He et al. 2021). TATA-box, CAAT-box, CGTCA-and TGACG-motif, ABRE, G-box, and MYB are the common *cis*-elements found among all the NR promoters (Fig. S1). Based on the available reference sequences, Maximum and least no. of *cis*-elements were found in *NR 6-1(A*, *B, D)* and *NR 7D* respectively.

**Fig. 4.**
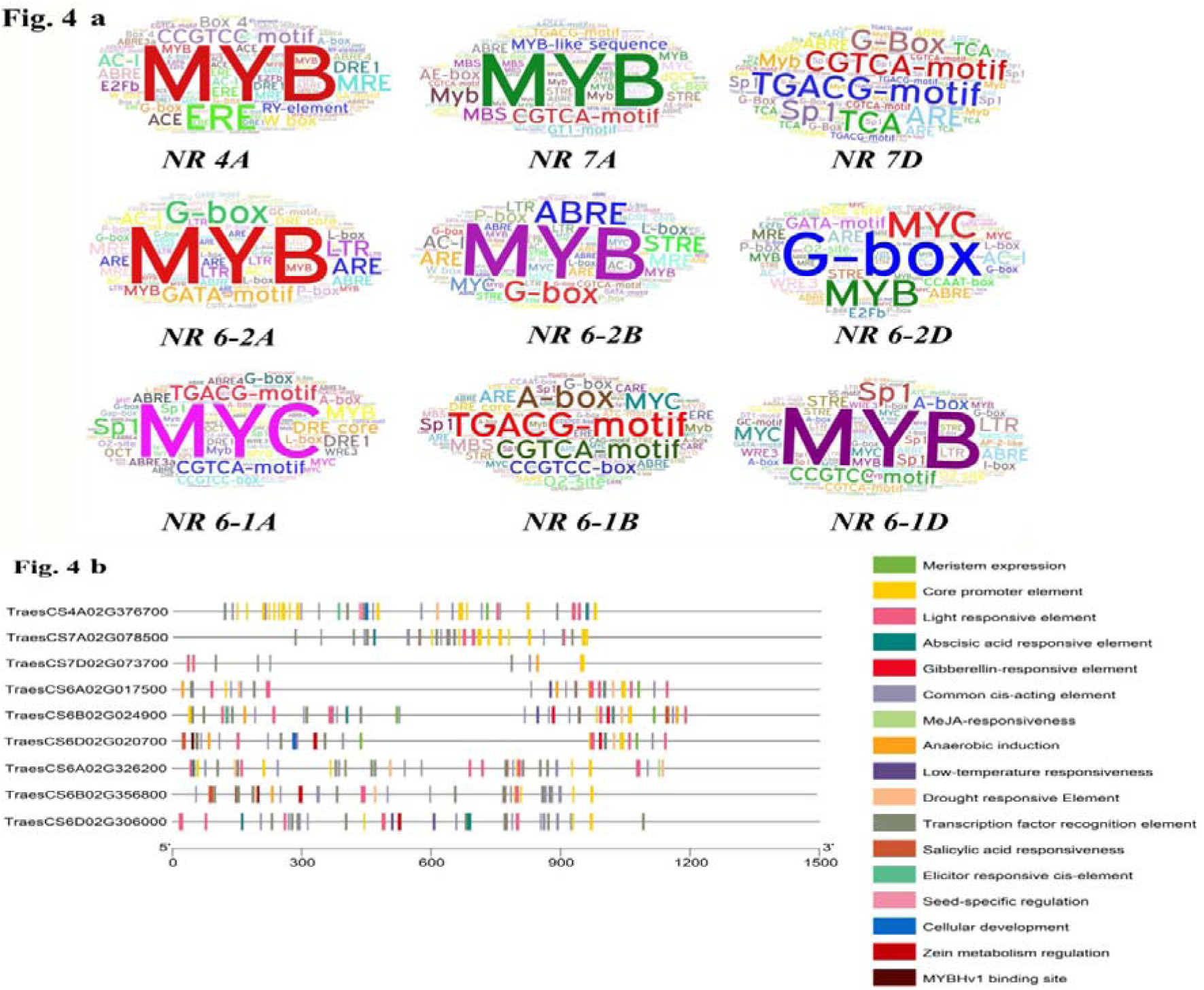
**(a)** The abundance of *cis*-acting elements (Except CAAT and TATA Box) in the promoter regions of *NRs* using the WordArt program. Font size increases with the abundance of *cis*-acting regulatory elements, **(b)** Distribution of *cis*-acting elements in promoter regions of *NRs*. Different colors represent different *cis*-acting elements. The scale bar is shown at the bottom

### Assessment of NUE related field parameters for selection of contrasting genotypes

Field parameters of HD 2967 and Choti Lerma genotypes were measured and are presented here. The GY/P of both the genotypes was significantly different under all the N levels during both the years. Whereas N uptake of both the genotypes was not significantly different at Y2-0 and Y1-0 (Fig. 5). The GY/P and N uptake was significantly higher in genotype HD 2967compared to Choti Lerma. Other parameters i.e. Biomass, Grain N Y/P, N at Harvesting, NUE (GY/P)/[(Soil N+Applied N)/P)] are presented in supplementary file (File S2).

**Fig. 5.**
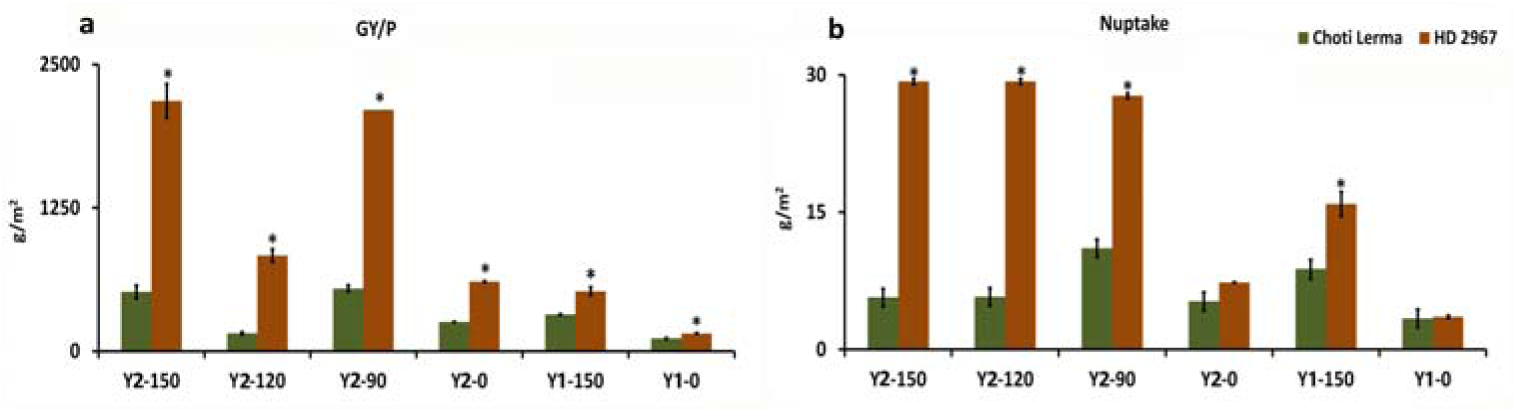
GY/P (Grain yield/plot) and N uptake of Choti Lerma and HD 2967 under different levels of N treatment in field condition. Y1 and Y2 denote 1^st^ and 2^nd^ year respectively. 150, 120, 90, 0 are the quantities (kg/ha) of N fertilizer applied. * denotes the significant difference among both the genotypes

### Performance of contrasting HD 2967 and Choti Lerma genotypes in hydroponics

There are significant differences in the shoot and root biomass parameters (Fig. 6) among both genotypes under both N-optimum and N-stress conditions except RDW under N-optimum and SDW under N-stress. All root and shoot parameters except RDW were significantly higher in HD 2967 under N-optimum condition. While under N-stress conditions, only root parameters i.e. RL, RFW and RDW were significantly higher in Choti Lerma (Fig. 6d, e, f). Chlorophyll (Chla, Chlb and Tchl) and carotenoids contents (Fig. 7) were significantly reduced in both genotypes under N-stress and these pigments were significantly higher in HD 2967 under both the conditions.

**Fig. 6.**
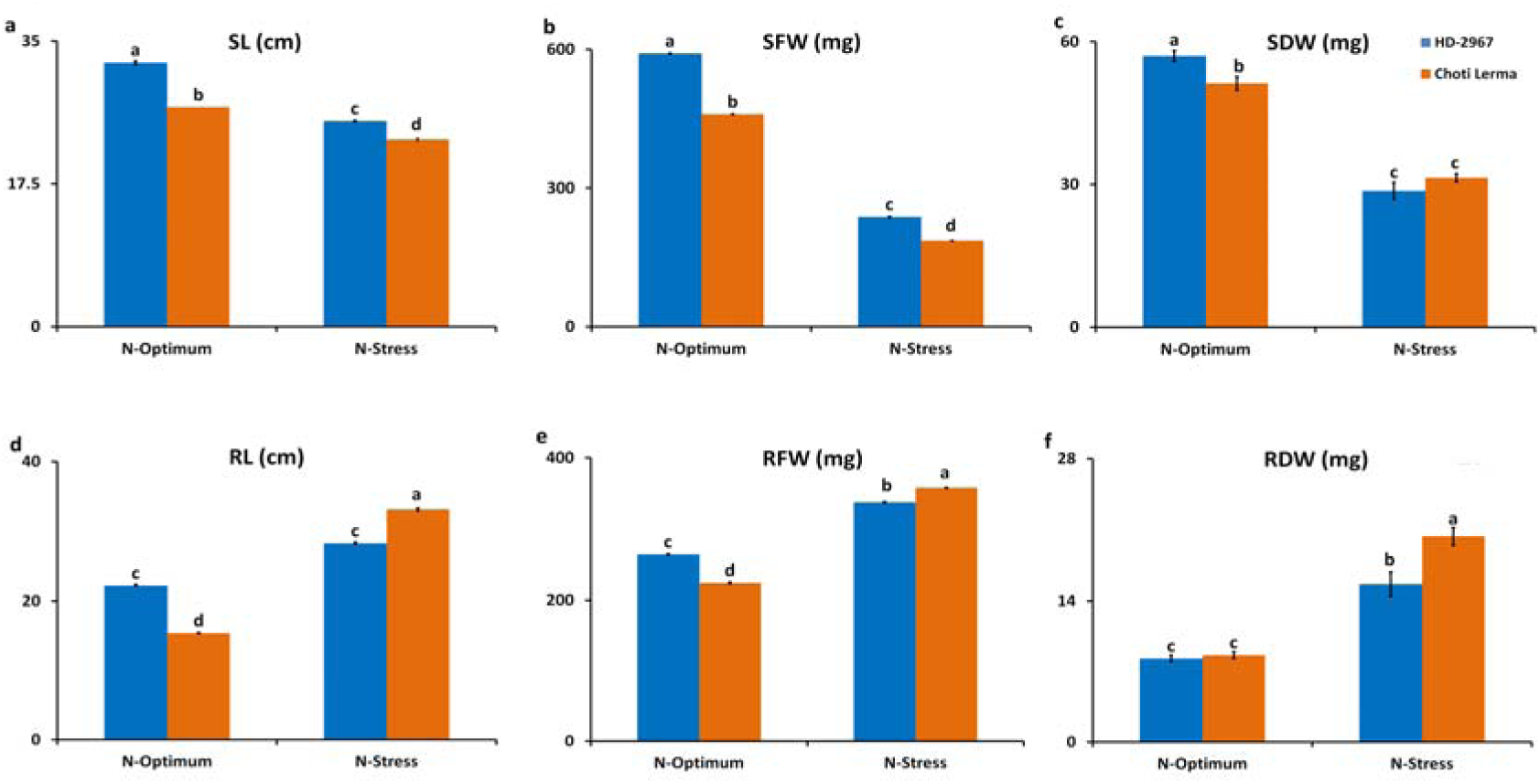
Shoot and root biomass parameters of 21 days old seedlings of HD 2967 and Choti Lerma under N-optimum and N-stress conditions. a) SL: Shoot length; b) SFW: Shoot fresh weight; c) SDW: Shoot dry weight; d) RL: Root length: e) RFW: Root fresh weight; f) RDW: Root dry weight

**Fig. 7.**
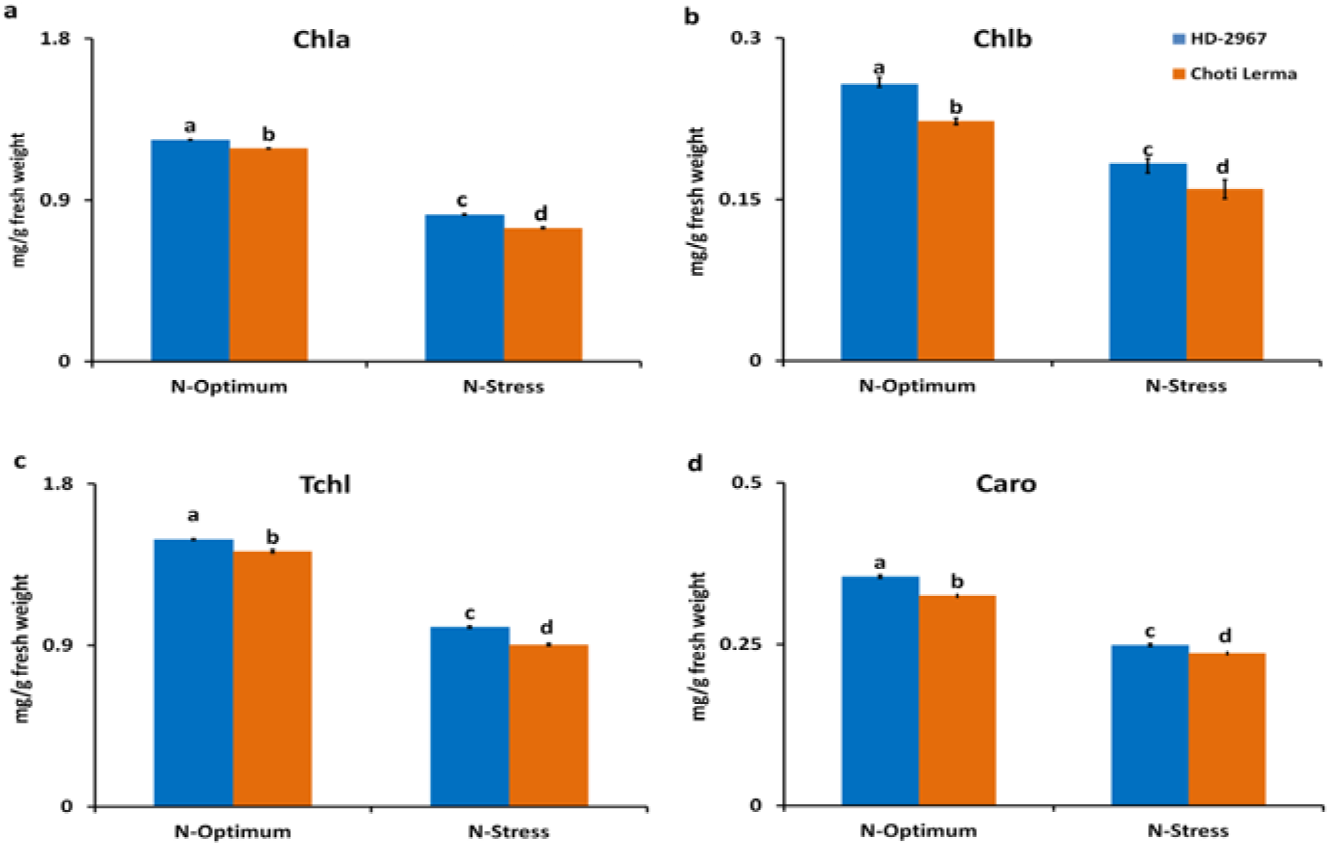
Chlorophyll and carotenoid contents of 21 days old seedlings of HD 2967 and Choti Lerma under N-optimum and N-stress conditions. a) Chla: Chlorophyll a content; b) Chlb: Chlorophyll b content; c) TChl: Total Chlorophyll content; d) Caro: Carotenoid content

Nitrogen content in shoots as well roots was measured under both N-optimum and N-stress conditions (Fig. 8). N-stress significantly reduced the SNC (Fig. 8a) while significantly increased the RNC (Fig. 8b) in both genotypes. SNC was found higher in HD 2967 under N-optimum condition while under N-stress condition both genotypes showed the similar level of nitrogen content. Contrastingly, RNC was found to be at similar level in both genotypes under N-optimum condition and found higher in Choti Lerma under N-stress condition. All over, TNC was found significantly different in both genotypes under both conditions but followed same trend as mentioned earlier (higher value in HD-2967 under N-optimum and higher value in Choti Lerma under N-stress condition) (Fig. 8c).

**Fig. 8.**
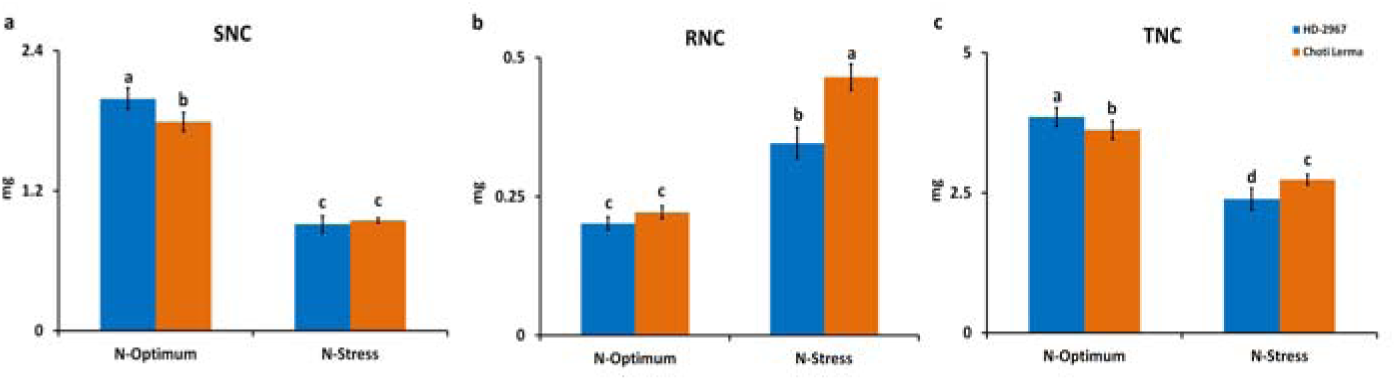
Nitrogen content of 21 days old seedlings of HD 2967 and Choti Lerma under N-optimum and N-stress conditions. a) SNC: Shoot nitrogen content; b) RNC: Root nitrogen content; c) TNC: Total nitrogen content

### NR specific activity in contrasting HD 2967 and Choti Lerma genotypes

NR activity was measured in the shoot tissues of 21 days old seedlings of HD 2967 and Choti Lerma under N-optimum and N-stress conditions. Results showed reduced NR activity under N-stress in both genotypes (Fig. 9). However, though the NR activity of HD 2967 was higher under N-optimum condition, Choti Lerma displayed a higher NR activity under N stress condition.

**Fig. 9.**
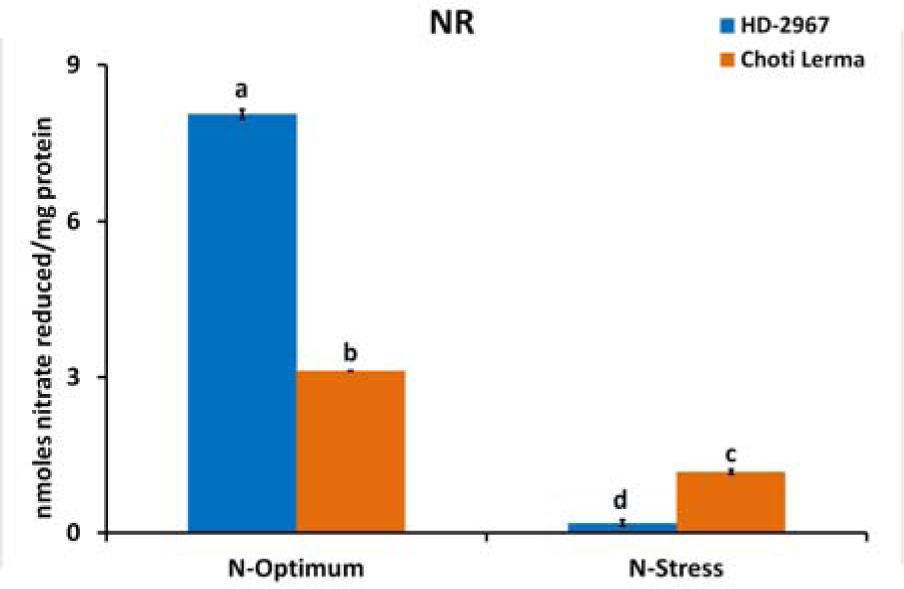
**NR specific enzyme activity shoot tissues of 21 days old seedlings of HD 2967 and Choti Lerma under N-optimum and N-stress conditions**

### Expression Pattern of *NR* genes under N-Stress by qRT-PCR Analysis

The expression pattern of three *NR* homeologue sets (*NR 6-1*, *NR 6-2,* and *NR 7AD-4A*) were studied by qPCR under Chronic N-stress and compared with N-optimum conditions. Expression studies revealed that the *NR 6-1* homeologue set has shown maximum responsiveness (maximum down-regulation) under N-stress (Fig. 10), which we took for our present study.

**Fig. 10.**
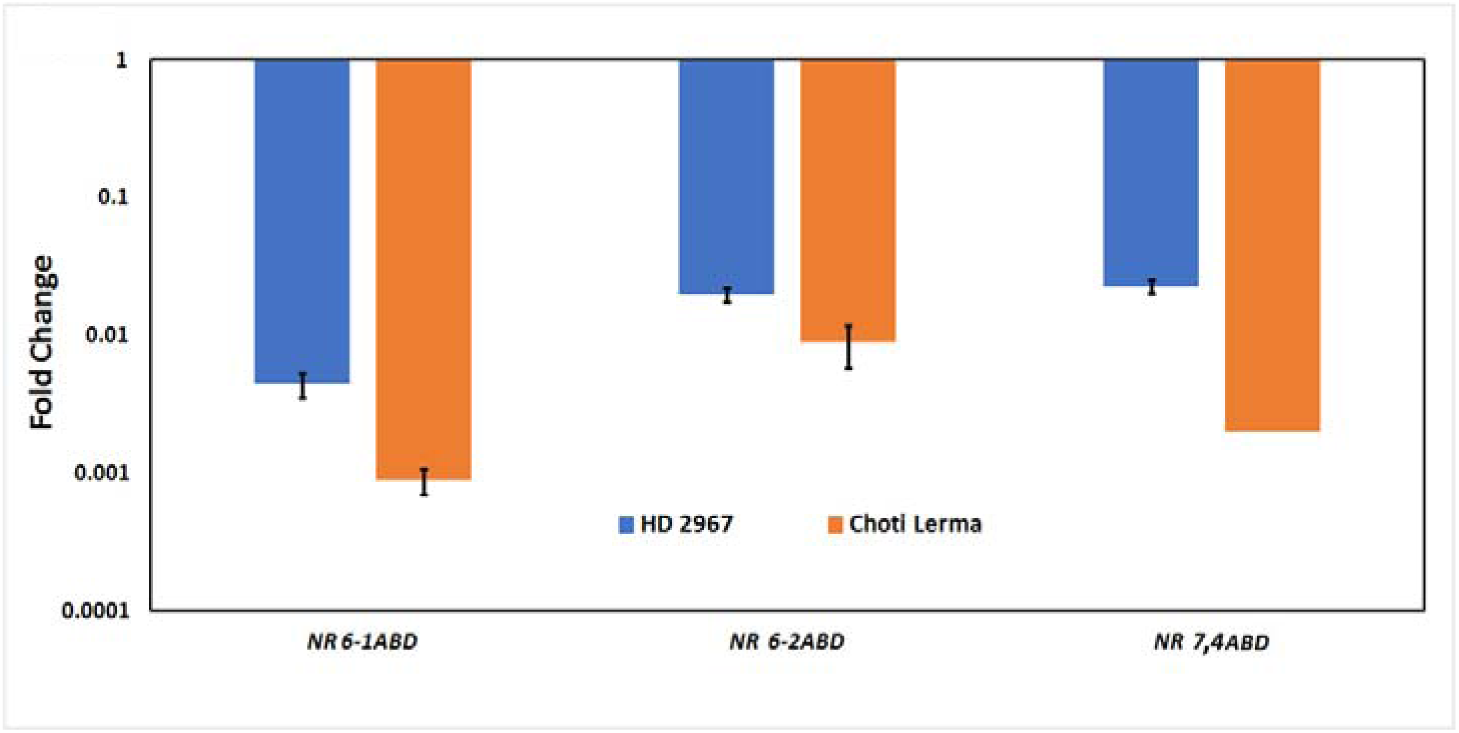
Relative expression of *NR 6-1, NR 6-2* and *NR 7, 4 ABD* homeologue sets in HD 2967 and Choti Lerma seedlings were grown under chronic N-stress conditions

### Homologue Variation Study of *NR 6-1A, NR 6-1B, NR 6-1D* genes from HD 2967 and Choti Lerma

The nucleotide sequences of cloned CDS of *NR 6-1A*, *NR 6-1B,* and *NR 6-1D* homeologues from both the wheat genotypes (HD 2967 and Choti Lerma) were confirmed using the BLAST tool in NCBI and then submitted to GenBank NCBI [NCBI accession numbers MT876649 (*NR 6-1A,* HD 2967), MT876650 (*NR 6-1B,* HD 2967), MT876651(*NR 6-1D,* HD 2967), MT876646 (*NR 6-1A,* Choti Lerma), MT876647(*NR 6-1B,* Choti Lerma), MT876648 (*NR 6-1D,* Choti Lerma). MSA (Fig. S2) revealed the homeologous differences in wheat for *NR 6-1* gene. ORF finder analysis of these sequences indicated 2926, 2874, and 2865 bp CDS for *NR 6-1A*, *NR 6-1B,* and *NR 6-1D* homeologue respectively, encoding 899, 896, and 895 amino acids that exhibited over 93% similarity to one another in both the genotypes. The polymorphisms were observed randomly over the entire CDS (Table S2 and Table S3). There were a total of 128 common SNPs (in both genotypes) found among the CDS of *A*, *B,* and *D* homeologues, of which 47 were transition and 81 were transversion, lead to a change in 32 amino acids. Other than SNPs, *B* and *D* homeologues of both genotypes were differentiated by 9 and 14 deletions in their respective CDS.

When the *NR* CDS of two contrasting genotypes were aligned, *NR 6-1A* showed 5 SNPs, of which 1 was transition and 4 were transversion, leading to a change in 3 amino acids; *NR 6-1B* showed 28 SNPs, of which 7 were transition and 21 were transversion, lead to a change in 24 amino acids and *NR 6-1D* showed 8 SNPs, of which 5 were transition and 3 were transversion, lead to a change in 5 amino acids (Table S4). Position of the nucleotides was counted from ATG, with A’s position as 1; similarly position of amino acids are counted with first amino acid M’s position as 1. SIFT analysis was done among the six amino acid sequences of NR 6-1 protein from the three homeologues each from HD 2967 and Choti Lerma. We found five conserved domains 1) oxidoreductase molybdopterin binding domain, 2) Mo-co (molybdenum cofactor) oxidoreductase dimerization domain, 3) a cytochrome b-5 like heme-binding domain 4) oxidoreductase FAD-binding domain 5) oxidoreductase NAD-binding domain. The results showed that the amino acid changes were found to be tolerated in all the homeologues of both the genotypes (Fig. 11).

**Fig. 11.**
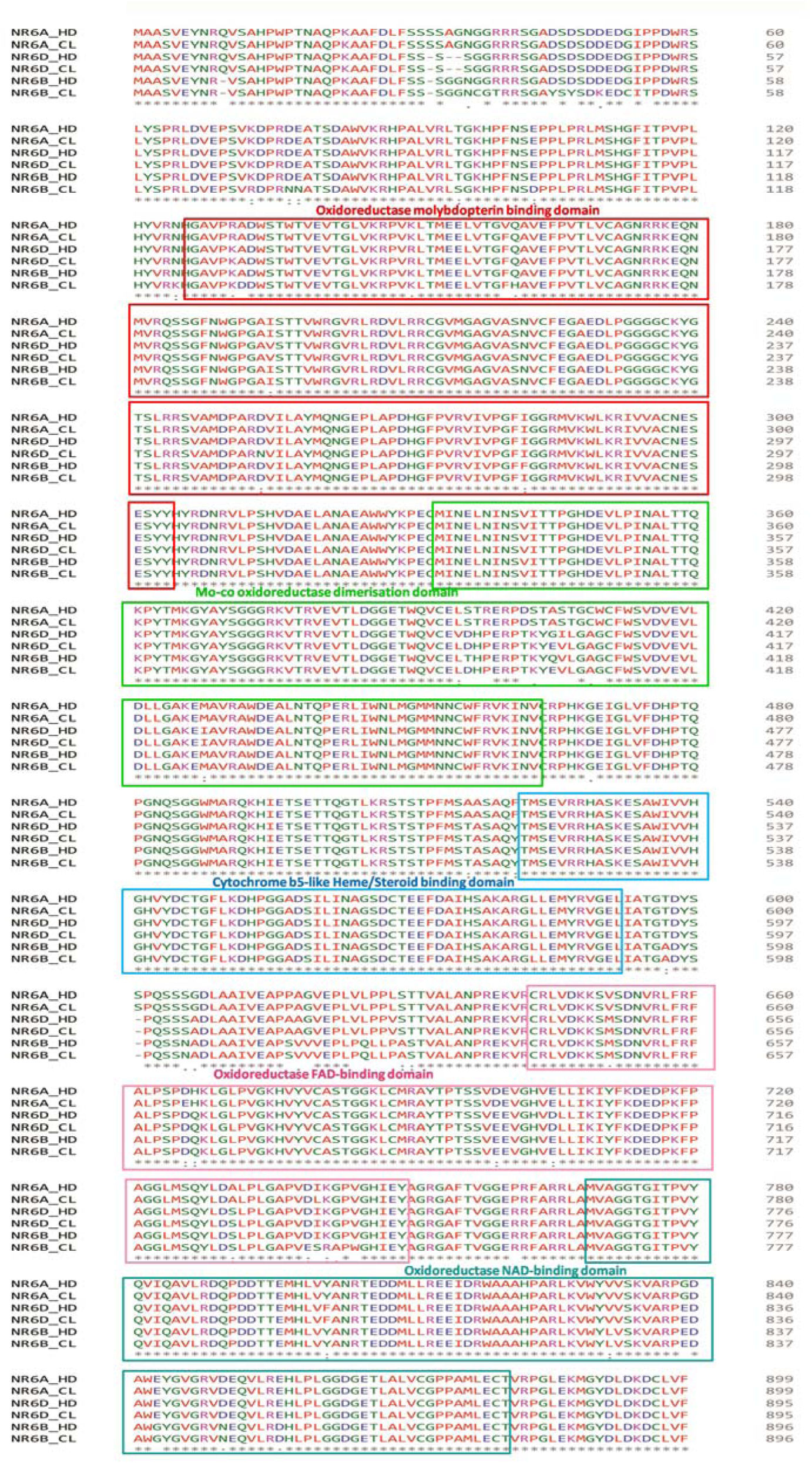
Changes in Functional Domains due to change in amino acids in homeologues of NR 6-1 protein in two contrasting genotypes, HD (HD 2967) and CL (Choti Lerma)

### Homeologue-specific expression profiles of *NR 6-1A, 6-1B, and 6-1D* genes under transient and chronic N-Stress

Results of qPCR confirmed that during N-stress, *NR* gene expression was downregulated under chronic N-stress in both the genotypes. When *6-1D-*homeologue was found most responsive in both the genotypes, the expression of *6-1A* homeologue was higher than *6-1B* homeologue in the case of HD 2967, but almost unchanged in the case of Choti Lerma (Fig. 12a).

**Fig. 12.**
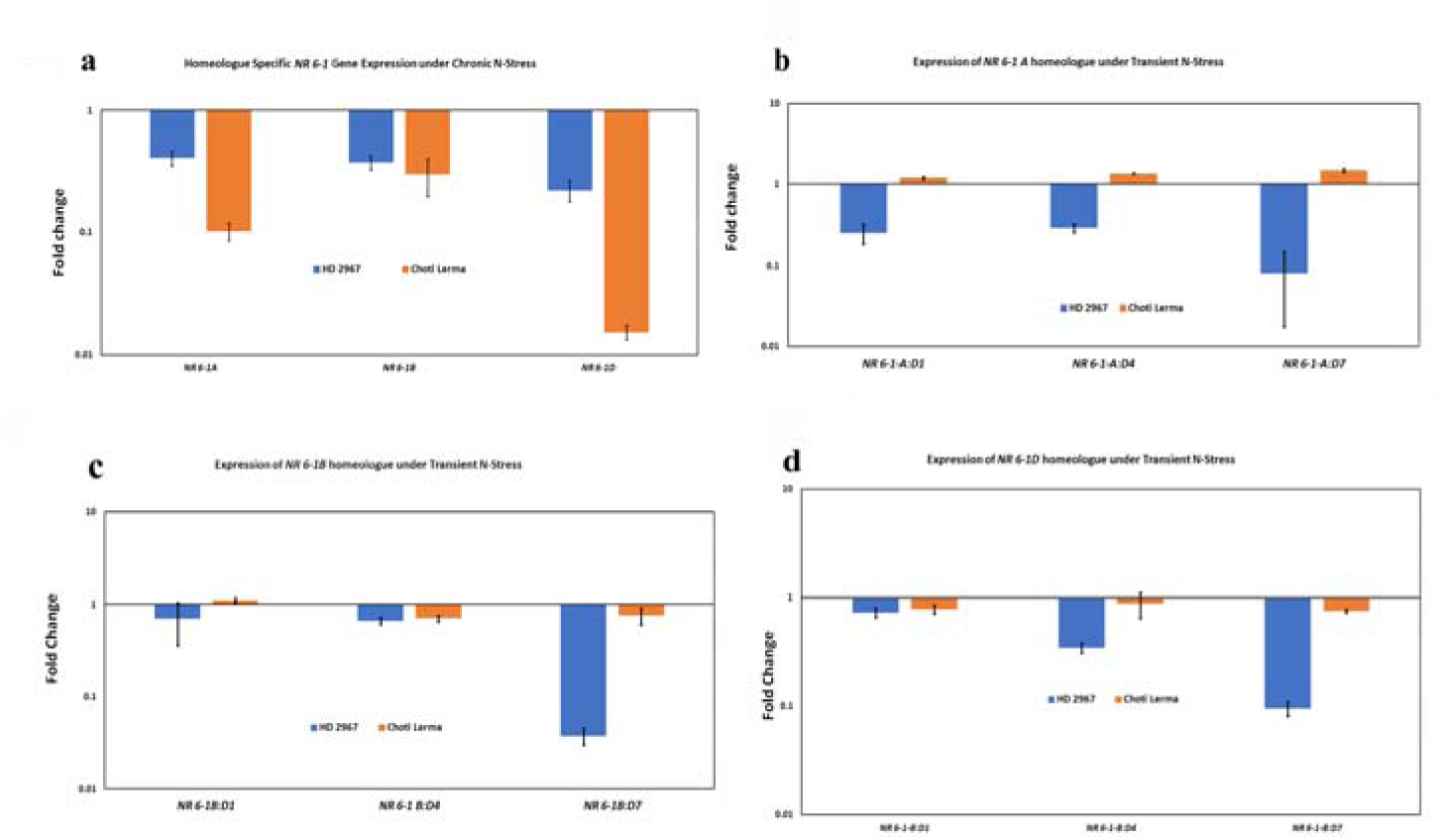
**(a)** Homeologue-Specific *NR 6-1(A, B, D)* Gene Expression Under Chronic N-Stress both the genotypes HD 2967 and Choti Lerma; **(b)** Expression of *NR 6-1A* homeologue under Transient N-Stress **(c)** Expression of *NR 6-1B* homeologue under Transient N-Stress; **(d)** Expression of *NR6-1D* homeologue under Transient N-Stress; 1D, 4D and 7D represent shoot tissues collected on 1 day (15 days After Transfer (DAT)), 4 days (17 DAT), and 7 (21 DAT) days after the imposition of N-stress

When the gene expression at the different time points of transient stresses was analysed, it was found that the expression of *NR 6-1A* homeologue was gradually downregulated in the case of HD 2967 over 10 fold during 7 days stress period whereas in the case of Choti Lerma, it was upregulated marginally (approximately 1.5 fold) gradually during the stress period (Fig. 12b).

The trend was almost similar for the other homeologues, *NR 6-1B* and *NR 6-1D* also, where the expression was gradually downregulated during 1 to 7 days of stress and it reduced to >30 and >10 fold respectively in the case of HD 2967 genotype (Fig. 8c and d). Again the changes were negligible in the case of Choti Lerma, though the trend was towards downregulation (Fig. 12c and d).

### Contribution of homeologues in HD 2967 and Choti Lerma

The contribution of each homeologue was also estimated in control, transient as well as in chronic stress conditions. In both the genotypes, the contribution of *NR 6-1B* homeologue was substantially less in comparison to *NR 6-1A* and *NR6-1D* homeologues in all the conditions. During transient stress, the contribution of *NR 6-1A* and *NR 6-1B* homeologue were gradually decreasing from 1 to 7 days of stress in the case of HD 2967, though they were statistically nonsignificant. However, it increased significantly during 1 to 7 days of stress in the case of *NR 6-1D* homeologue. Though *6-1B* homeologue had the least contribution, under N-optimum and N-Chronic stress conditions, its contribution was higher than transient stress up to 7 days (Fig. 13a) in HD 2967 genotype.

**Fig. 13.**
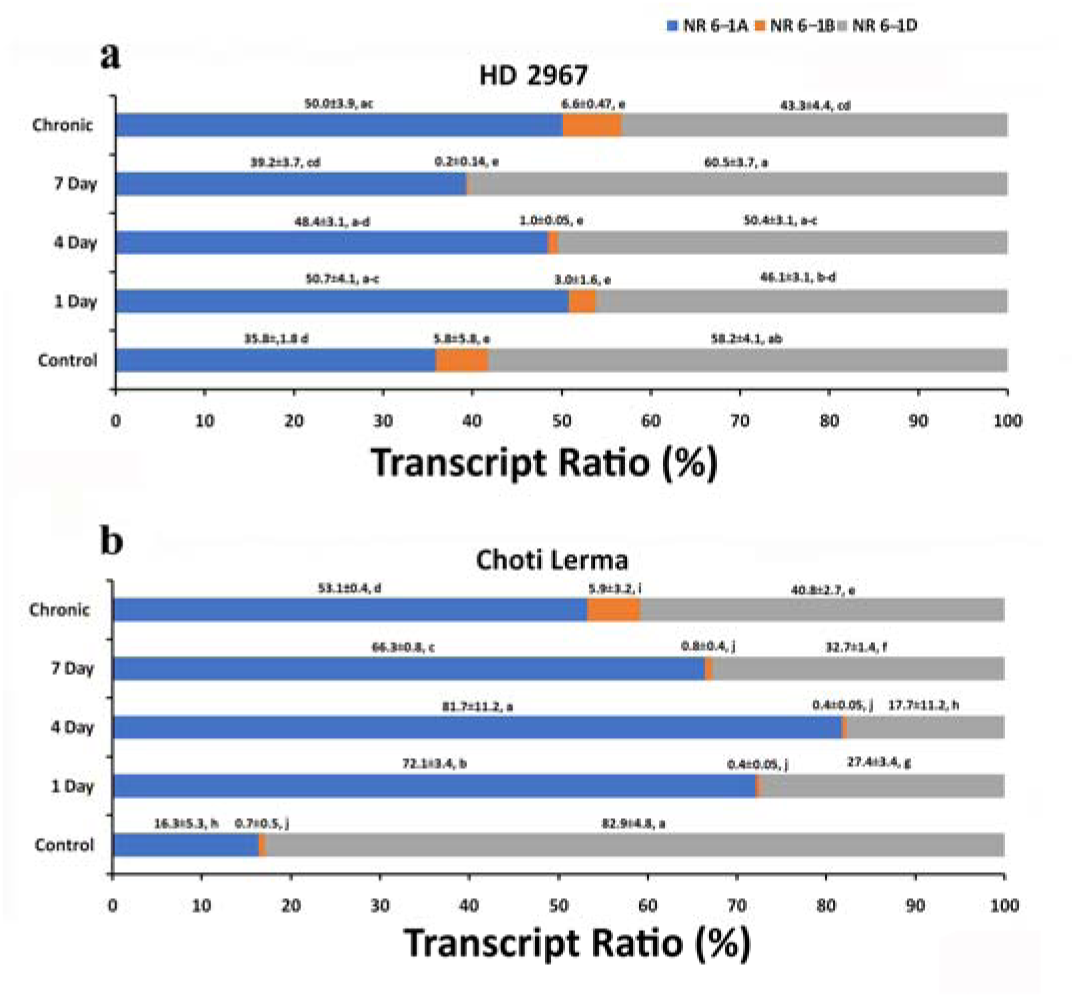
**(a)** Contribution of each *NR 6-1* homeologues in HD 2967 genotype under N-optimum and different N-Stress conditions **(b)** Contribution of each *NR 6-1* homeologues in Choti Lerma genotype under N-optimum and different N-Stress conditions

In Choti Lerma, the contribution of *NR 6-1B* in chronic stress was significantly higher than the rest of the conditions including N-optimum. The contribution of *NR 6-1A* homeologue increased to a significantly higher level during chronic as well as transient stress. The contribution of *NR 6-1D*-homeologue is higher in the case of the N-optimum condition (Fig. 13b).

## Discussion

Most of the crops including wheat, nitrate is the preferred form of nitrogen in the aerobic soil, though plants can uptake other forms of nitrogen too including ammonia, amino acids, and peptides, which depends on the physio-chemical properties of the soil (Tsay et al. 1993; Krouk et al. 2006; Nour-Eldin et al. 2012). In normal growth conditions, nitrate is taken up from the soil and assimilated mostly in shoots to form glutamate, the first amino acid, which is subsequently used for N-containing biomolecules production. NUE depends on multiple factors, mainly on their uptake, translocation, and assimilation efficiency (Li et al. 2017). However, it is not necessary that efficient uptake and translocation will lead to efficient assimilation. Hence, in this study, we focused on assimilation, especially on the first step of assimilation. As mentioned earlier, after nitrate is taken up by the plant from the soil, the assimilation process starts by NR and is reduced to nitrite by this important enzyme. It has also been reported that the *OsNR2* allele of the *NR* gene can increase the nitrogen use efficiency in rice (Gao et al. 2019). It is well reported that this enzyme is responsive to nitrogen stress, mainly nitrate stress in different cereal crops including wheat (Botella et al. 1993; Golberg et al. 1995)(Botella et al. 1993; Golberg et al. 1995; Nathawat et al. 2005; Santos et al. 2019). Though an extensive study on *NR* gene has been conducted in *Arabidopsis* and Rice, understanding of this gene in wheat is limited. Two *NR* genes in *Arabidopsis* (Wilkinson and Crawford 1991) and three in Rice (Hamat et al. 1989) have been identified and most of them have been well characterized.

In our study, we also found nine copies of the *NR* gene in hexaploid bread wheat as reported recently by Hurali et al. 2022 (Hurali et al. 2022). Since there are three genomes in this crop, three copies are present for most of the genes in the bread wheat genome (Juery et al. 2021), but that is not the case with *NR*. Two copies were found in the B genome, three in the D genome, and four copies in the A genome. We further categorized the *NR*s into three groups (clusters) with three copies in each cluster based on their sequence similarity analysis. Mostly we saw multiple copies of the genes for the essential functions of the organisms such as the *WOX* gene, 43 copies of the gene are present in wheat, 13 copies in rice, and 15 in *A. thaliana* (Zhang et al. 2010; Rathour et al. 2020) and that is because these genes are essential for the growth and development of plants. Likewise, in the present study, being the most essential and key enzyme for nitrogen metabolism, *NR* is also found in multiple copies in wheat. Though these copies possibly work as backup for the plant for carrying out the essential function of nitrate assimilation, these many copies of *NR* may look a little too much. However, reports also say that gene diversification is common in plants, and in many cases, the diverse genes get changed to other functions also (Panchy et al. 2016). In bread wheat, the gene diversification might not have happened solely at hexaploid level; but at the diploid or tetraploid (progenitors) level. Hence, the number of copies of a gene present in bread wheat is the combination of pre-and post-polyploidization. Among the nine copies of the *NR* genes, diversification of the copies was found among these three sets in terms of gene structure, predicted protein size and motif configuration as mentioned in the recent research (Hurali et al. 2022), which suggest similar function of the copies within a set, as also reported for other proteins in wheat (Gao et al. 2021). Collinearity analysis in our study suggested that the increase in *NR* copies and their divergence in hexaploid wheat might be due to the polyploidy events (Fig. 3), where diploids have 2-3 copies, tetraploid has 6 and finally hexaploid has 9 copies of the gene. In spite of having relatively large size (5-8 kb), all the copies of *NR* gene in wheat had only a few introns (1 or 2), which is rare in case of other genes of similar size. However, this feature is not *T. aestivum* specific, but *NR* gene specific, which we found also in case of tomato, rice and tobacco (Choi et al. 1989; Daniel-Vedele et al. 1989; Vaucheret et al. 1989). Out of the nine *NR* copies, six copies have two and three copies have one intron. This loss of one intron might be due to occurrence of events over evolutionary time (Wang et al. 2017). Hurali et al. (2022) (Hurali et al. 2022) attempted to derive a phylogenetic relation of these nine copies of *NR* involving only three cereal species (wheat, rice, maize) and Arabidopsis. We extended the study involving 22 different monocot and 83 dicot species. As expected, our study revealed the closest similarity of hexaploid bread wheat with tetraploid durum wheat followed by the diploid progenitors, and then with *H. vulgare*. However, interestingly, the NR does not form monocot and dicot specific separate groups, unlike other N-assimilating enzymes such as CS, ICDH, GS2, Fd-GOGAT (Gayatri et al. 2019, 2021). Hence, it appears that the amino acid sequences of NR proteins have been more conserved during the evolution of the NR gene (Campbell 1996; Zhou and Kleinhofs 1996).

NR enzyme activity as well as gene expression are known to be reduced under N-stress in wheat (Chandna et al. 2012; Zhao et al. 2013; Kavoosi et al. 2014; Sinha et al. 2015). Analysis of *cis-*element shows the prominent presence of MYB in almost all the copies of *NR* genes (Fig. 4), which is a known N-responsive element in other plants (Lea et al. 2007; Konishi and Yanagisawa 2010). In addition, presence of MYC, G-Box, GATA-motif is also reported to be N-responsive in different plants (Witko 2003; Konishi and Yanagisawa 2013; Zhang et al. 2015; He et al. 2021; Islam et al. 2021). Though there is no report on N-responsive elements in wheat genes, this analysis confirms N-responsive of the above elements in wheat. However, the list of these N-responsive elements in *NR* gene may not be complete and comprehensive, as there are still gaps in the reference genome sequence. Presence of these N-responsive elements are occurring in combination with the phytohormone-responsive elements in the promoter region indicates the complex regulatory network between the N assimilation and hormonal pathways (He et al. 2021).

Gene expression of the three sets of *NR* was downregulated under N-stress, which was expected (Gayatri et al. 2019, 2021), but the most responsive was *NR 6-1ABD*. Further, we studied this *NR 6-1ABD* in detail considering the three homeologues in two contrasting wheat genotypes HD 2967 and Choti Lerma. It is important to mention here that the contrasting nature of these two genotypes was established using the field NUE related parameters. GY/P and N-uptake were significantly higher in HD 2967 compared to Choti Lerma measured during the two consecutive years at all doses of applied N, except N-uptake @ 0kg/ha applied N. There were a few reports on the nature of these two genotypes (Chattaraj et al. 2013; Nagar and Gayatri 2018; Paul 2018; Tyagi et al. 2020) but without enough supporting data. HD 2967 generally exhibited superior performance in terms of biomass parameters, pigment content, nitrogen content, and NR activity under N-optimum conditions. However, under N-stress conditions, Choti Lerma showed an advantage in root parameters and NR activity. The higher biomass parameters observed in HD 2967 under N-optimum conditions suggest its ability to efficiently utilize nitrogen resources for growth. Similarly, the increased pigment content in HD 2967 indicates a better capacity for photosynthesis, potentially leading to enhanced plant performance. Moreover, the higher nitrogen content and NR activity in HD 2967 under N-optimum conditions further emphasize its better nitrogen responsiveness. Two-way comparisons were made using the sequences of each homeologues of the two contrasting genotypes: one way was to study allelic variation among *NR 6-1A, NR 6-1B,* and *NR 6-1D* homeologues; and the other way was the allelic variation of each homeologue between the two genotypes (Fig. 14).

**Fig. 14.**
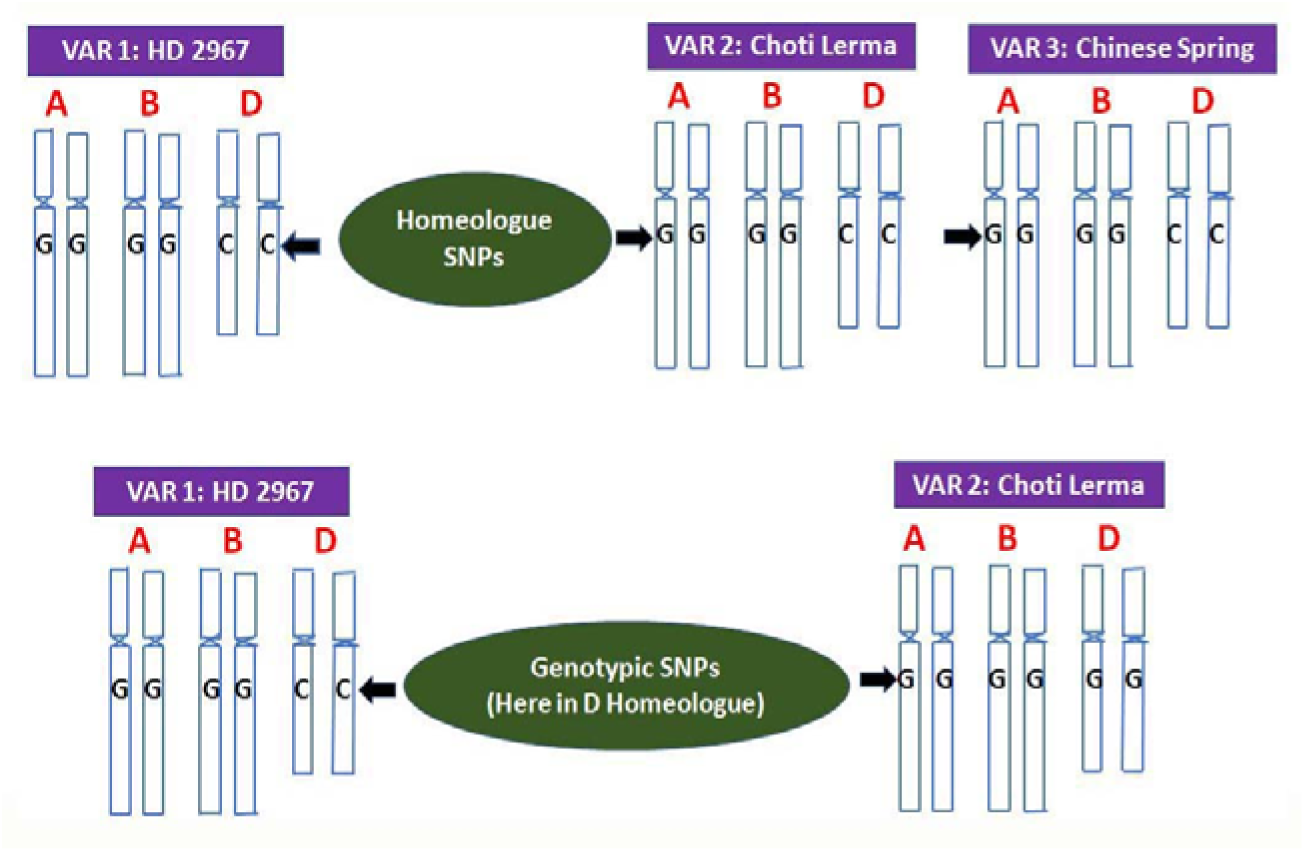
Top panel shows the variation among the homeologues where the gene in *D* homeologue is different from *A* and *B* homeologues and has SNP; the Bottom: shows the genotypic difference among the homeologues where the gene in *D* homeologue is having SNP

When each homeologues were compared between HD 2967 and Choti Lerma, the nucleotide polymorphism was very low with proportional variation in amino acid changes. Since the homeologues have been derived from three different species, the polymorphism among the homeologues was expected to be more than that of the genotypic differences of an individual homeologue (Fa et al. 2010; Winfield et al. 2012) and we found the expected result. However, interestingly the transversion was significantly more than transition which is contrary to the general observations for any genes (Winfield et al. 2012; Alipour et al. 2017; Rimbert et al. 2018; Zhou et al. 2020). Probably this phenomena of higher transversions is *NR 6-1* specific, which has led to more number of amino acid changes, though they were tolerated in the functional domains of the enzyme and functionality was retained. However, such a higher rate of transversion in the gene does exist, e.g, *CD14* gene *in Bos taurus* and *Bos indicus* (Morenikeji et al. 2020); *MYB* gene in chickpea (Roorkiwal et al. 2014) etc.

We also studied the homeologous gene expression under chronic as well as transient stresses. Under the optimal condition, nitrate assimilation in the shoot is more than that of the root, and hence, studying gene expression in the shoot was important. Though we found the presence of common *cis*-elements in the promoters of a set of homeologue, the proportion of the elements differs from one homeologue to the other (Fig. 4). The presence of N-responsive elements like MYB, MYC, GATA-motif and G-Box were almost missing in B homeologue, whereas MYC, MYB were very prominent in *A* and *D* homeologues respectively of *NR 6-1*. Contribution of the homeologues are corroborating with the presence of N-responsive elements in the homeologues, where B had a low relative expression under N-stress in comparison to A and D in case of *NR 6-1*. Homeologue-specific expression of *NR 6-1A, NR 6-1B,* and *NR 6-1D* in the two genotypes under chronic and transient stresses were conducted keeping a few questions in mind. 1. How does each homeologue of this specific *NR* gene respond to nitrogen stress in wheat? 2. How the transient stress alters the gene expression over a period of time? 3. How long-term stress from the beginning of growth (chronic stress) differs from the sudden stress during normal growth (transient)? 4. How the expression alters when the transient stress period is prolonged? 5. How do the two very contrasting genotypes with respect to N-responsiveness differ in gene expression, both under chronic and transient stresses? 6. What is the contribution of different homeologue of this gene? and finally, 7. Whether the contribution patterns are same in both the genotypes?

Answers of all the questions are revealed from the study, which we have discussed here. 1. As expected, each homeologues gets downregulated especially in case of chronic stress (Botella et al. 1993; Golberg et al. 1995; Santos et al. 2019; Gao et al. 2021), in fact, all the three sets of *NR* genes were downregulated under N-stress. However, the *NR 6-1D* homeologue was most affected in both the genotypes under chronic stress. 2. Transient stress and chronic stress differed the gene expression. Downregulation of *NR 6-1B* homeologue was maximum under stress, especially at 7 days after the imposition of transient stress. 3. When the stress period was imposed suddenly after normal growth, initially up to 3 days, the change in gene expression was not comparable to chronic stress, but after 7 days of transient stress, the effect was at par with the chronic stress in the case of HD 2967. 4. In the case of transient stress, the gene expression was gradually downregulated in all three homeologues, and lowest expression was at 7 days after the imposition of stress. 5. Choti Lerma, a less N-responsive genotype, was minimally affected with transient stress in comparison to HD 2967. However, the gene expression was considerably downregulated for Choti Lerma in case of chronic stress, though not as much as HD 2967. Results indicate that Choti Lerma has a lesser responsive *NR 6-1* gene, which might be one of the causes of its poor N-responsiveness and probably vice versa. Since the requirement of nitrogen is less for this genotype, its immediate response to N-stress after a period of normal growth is very less as observed by Chandna et al. 2012 (Chandna et al. 2012).

Regarding questions no. 6 and 7, a clear observation and conclusion could be made about very low contribution of *NR 6-1B* homeologue; whereas *NR 6-1A* and *D* shared the major contribution between them under different conditions. We could also notice that contribution of *NR 6-1A* homeologue increased and *NR 6-1D* homeologue decreased significantly in the case of Choti Lerma under any type of N-stress compared to HD 2967. It was reported earlier while analyzing the homeologous contribution of *Citrate Synthase* (*CS*) and *Isocitrate dehydrogenase* (*ICDH*), genes of two vital carbon metabolizing enzymes linked with N-assimilation, that the *B-* homeologue was contributing least in the case of *CS* gene under N-optimum as well as N-stress condition, whereas the contribution was maximum for same homeologue in case of *ICDH* (Gayatri et al. 2019). The contribution of the homeologous genes of hexaploid bread wheat has been reported from the last decades onwards, even when the wheat genome sequencing project did not launch (Roorkiwal et al. 2014). Bias expression of the homeologues are also reported such as *dihydroflavonol-4-reductase* in root, leaves and grains (Himi and Noda 2004), *benzoxazinone (Bx*) biosynthetic genes (Nomura et al. 2005), *MADS-box* gene (Shitsukawa et al. 2007), etc., all of them showed a differential expression of *B*-homeologue. However, precise role of each homeologue is not yet deciphered, which is only possible if we mutate the other copies and homeologues of a gene, as suggested by Garcia-Oliveira et al. 2013; Tovkach et al. 2013 (Garcia-Oliveira et al. 2013; Tovkach et al. 2013). Probably the least contributing *NR 6-1A* gene would play a major role as back up when other copies are defective (mutated). However, combining the result of our study and the previous knowledge, it could be said clearly that bread wheat has a strong backup for most of the important genes with multiple copies. Contribution of a gene homeologue is condition specific and regulated by the promoter elements. Homeologous expression of any gene in wheat is of great interest (Shitsukawa et al. 2007) and the information from the present study will add up to the knowledge of the *NR* gene, especially in wheat crop and will be useful for its manipulation in the future.

## Conclusions

In this study, a total of 9 copies of the *NR* gene were categorized into three groups *NR 6-1ABD, NR 6-2ABD,* and *NR 7AD-4A. Cis*-element analysis also revealed that the genes are regulated by their N-responsive elements like MYB, MYC, G-Box and GATA-motif. Further, among the three sets, *NR 6-1 ABD* was found most responsive under N-stress. Cloning and sequence analysis of three homeologous CDS of *NR-6-1* from two contrasting wheat genotypes HD 2967 and Choti Lerma genotypes revealed that the homeologous difference in gene sequence was much more than the genotypic difference, and unusually transversion was more than a transition in the case of SNPs (Fig. 15). *D*-homeologue was found to be most responsive (affected and downregulated) to N-stress in both the genotypes. N-stress showed a quicker response in case of HD 2967 in comparison to Choti Lerma during 1 to 7 days period of stress. Comparatively lesser response of *NR 6-1* to N-stress in case of Choti Lerma is probably due to low N-requirement of this genotype. We found relatively very less contribution of *B*-homeologue, which was linked to the inconspicuous presence of N-responsive promoter elements (Fig. 11); however, the contribution of this homeologue under chronic stress was significantly higher than that of transient stress. We also found the expression of *A*-homeologue increased to a significantly higher level under N stress. This study is expected to help in manipulating the specific homeologue of the *NR* gene for desired assimilation of N in wheat. To continue with this work, *6-2 ABD* and *7AD-4A NR* homeologues should also be characterized in detail.

**Fig. 15.**
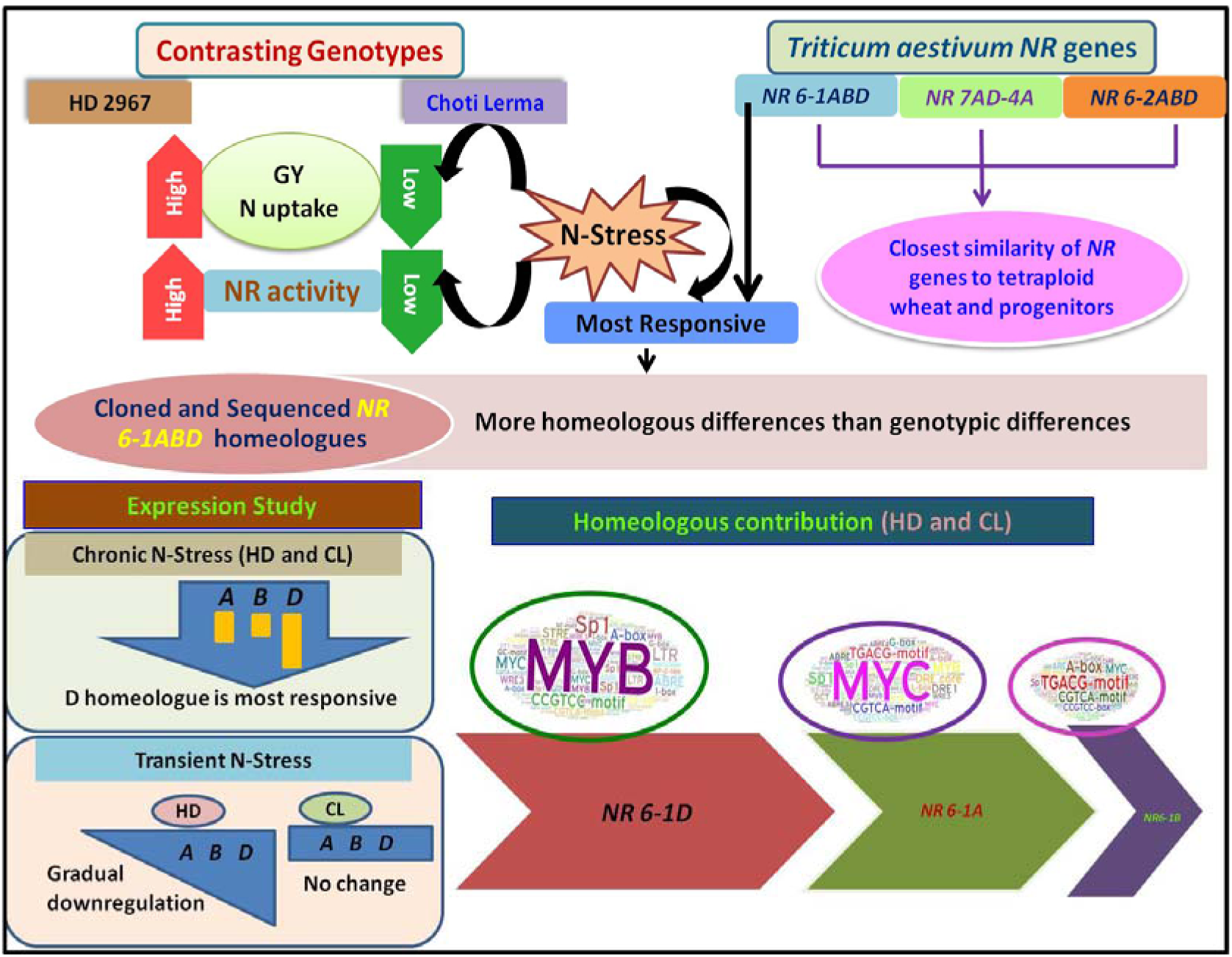
Summary of Identification and characterization of the *NR* gene in wheat (*Triticum aestivum* L.) and expression pattern along with homeologous contribution of *NR 6-1ABD* under N-stress in HD 2967 (HD) and Choti Lerma (CL). 9 *NR* genes were identified from wheat genome and grouped in to three sets. Phylogenetic analysis of these 9 wheat *NR* genes with other *NRs* from different monocot and dicot species displayed their closest similarity with diploid progenitors and tetraploid wheat. Among the three sets, *NR 6-1ABD* was found most responsive under N-stress. Cloning of three homeologous (*NR 6-1A, NR 6-1B, NR 6-1D)* CDS in two contrasting genotypes (HD and CL) revealed homeologous differences more than genotypic differences. Under chronic N-stress among the three homeologues, *D*-homeologue was found to be most responsive. Under transient N-stress, all the homeologues were gradually downregulated in the case of HD, whereas, in case of CL, there was found no significant change in the expression of these homeologues. Very less contribution of *B-* homeologue was found which can be linked to the absence of some N-responsive promoter elememts.

## Declarations

### Data availability statement

The CDS sequences of the genes were submitted to GenBank NCBI and the corresponding gene accession numbers are provided in the text and can be obtained in the NCBI nucleotide Database (https://www.ncbi.nlm.nih.gov/nucleotide/). All data generated or analyzed during the present study are included in this manuscript and its supplementary information files.

### Authors’ Contributions

Gayatri and Megavath Ravi designed and performed the experiments. Gayatri wrote the manuscript. Harsh Chauhan performed the hydroponics experiment. Ekta Mulani constructed circos. Sachin Phogat edited the manuscript. Karnam Venkatesh provided seeds of HD 2967 and Choti Lerma genotypes and all relevant data generated from the field conditions of both the genotypes, edited the manuscript. Pranab Kumar Mandal conceptualized the research objectives, designed the experiments, supervised the entire work, and revised the manuscript.

## Supporting information

Suppl.zip

## Acknowledgements

We would like to thank to the Department of Biotechnology, Government of India (BT/IN/UK-VNC/43/KV/2015-16) for funding under the project “Indo-UK Centre for the Improvement of Nitrogen Use Efficiency in Wheat (INEW)” and Director, NIPB, New Delhi for providing all the facilities. Authors also want to thank P.G. School, ICAR-Indian Agricultural Research Institute since part of the work is from Mr. Megavath Ravi’s M.Sc. thesis, submitted to this Institute.

## Abbreviations

CDS: Coding sequence
mmol/L: Millimolar/litre
h: Hour
cDNA: Complementary deoxy-ribonucleic acid
dNTPs: Deoxynucleoside triphosphates
PCR: Polymerase chain reaction
qPCR: quantitative-Polymerase chain reaction
T_m_: Melting temperature
°C: Degree Celsius
ORF: Open reading frame
MW: Molecular weight
pI: Isoelectric point
NCBI: National Center for Biotechnology Information
µl: Microlitre
ng: Nanogram
bp: Basepair
UTR: Untranslated region

