## Supplementary material for "Characterization and Expression Analysis of *Nitrate Reductase 6-1ABD* Gene in Hexaploid Bread Wheat Under Different Nitrogen Regime": Suppl.zip: Table S2.docx

**Table1 S2 Changes in amino acid in NR 6-1 protein due to SNPs among *A*, *B* and *D* homeologues of *NR 6-1* in wheat**

| **Sl. No.** | **Position** | ***NR 6-1A*** | ***NR 6-1B*** | ***NR 6-1D*** | **Sl. No.** | **Position** | ***NR 6-1A*** | ***NR 6-1B*** | ***NR 6-1D*** | **Sl. No.** | **Position** | ***NR 6-1A*** | ***NR 6-1B*** | ***NR 6-1D*** |
| --- | --- | --- | --- | --- | --- | --- | --- | --- | --- | --- | --- | --- | --- | --- |
| **1** | **36** | **G** | **C** | **G** | **45** | **1447** | **C** | **G** | **C** | **89** | **1975** | **C** | **C** | **G** |
| **2** | **141** | **T** | **C** | **C** | **46** | **1507** | **G** | **C** | **G** | **90** | **1981** | **C** | **C** | **G** |
| **3** | **177** | **G** | **C** | **G** | **47** | **1550** | **G** | **A** | **A** | **91** | **1990** | **G** | **C** | **G** |
| **4** | **210** | **G** | **A** | **G** | **48** | **1552** | **C** | **C** | **G** | **92** | **2005** | **C** | **G** | **G** |
| **5** | **354** | **G** | **C** | **C** | **49** | **1555** | **C** | **C** | **G** | **93** | **2014** | **C** | **G** | **G** |
| **6** | **392** | **G** | **A** | **A** | **50** | **1558** | **C** | **C** | **A** | **94** | **2041** | **C** | **C** | **G** |
| **7** | **444** | **C** | **G** | **G** | **51** | **1566** | **T** | **A** | **A** | **95** | **2101** | **C** | **G** | **G** |
| **8** | **447** | **C** | **G** | **G** | **52** | **1594** | **G** | **C** | **G** | **96** | **2113** | **C** | **C** | **T** |
| **9** | **449** | **A** | **A** | **G** | **53** | **1645** | **C** | **G** | **G** | **97** | **2116** | **C** | **G** | **G** |
| **10** | **489** | **C** | **C** | **G** | **54** | **1669** | **C** | **T** | **C** | **98** | **2119** | **G** | **G** | **C** |
| **11** | **507** | **C** | **G** | **C** | **55** | **1693** | **C** | **C** | **T** | **99** | **2152** | **C** | **T** | **C** |
| **12** | **555** | **A** | **G** | **C** | **56** | **1756** | **C** | **C** | **T** | **100** | **2195** | **G** | **T** | **T** |
| **13** | **582** | **A** | **A** | **G** | **57** | **1793** | **A** | **G** | **A** | **101** | **2197** | **G** | **C** | **G** |
| **14** | **585** | **C** | **T** | **C** | **58** | **1807** | **T** | **G** | **G** | **102** | **2252** | **C** | **T** | **C** |
| **15** | **606** | **G** | **C** | **C** | **59** | **1810** | **G** | **A** | **G** | **103** | **2267** | **C** | **C** | **G** |
| **16** | **648** | **C** | **G** | **G** | **60** | **1821** | **G** | **A** | **G** | **104** | **2287** | **C** | **G** | **G** |
| **17** | **660** | **C** | **G** | **C** | **61** | **1824** | **G** | **C** | **C** | **105** | **2339** | **C** | **C** | **G** |
| **18** | **671** | **C** | **C** | **T** | **62** | **1849** | **C** | **T** | **C** | **106** | **2402** | **C** | **G** | **G** |
| **19** | **804** | **C** | **G** | **C** | **63** | **1852** | **T** | **C** | **T** | **107** | **2407** | **A** | **A** | **T** |
| **20** | **810** | **C** | **C** | **T** | **64** | **1853** | **C** | **T** | **G** | **108** | **2420** | **G** | **C** | **C** |
| **21** | **852** | **C** | **G** | **C** | **65** | **1857** | **C** | **T** | **C** | **109** | **2474** | **C** | **C** | **G** |
| **22** | **936** | **G** | **C** | **G** | **66** | **1860** | **G** | **T** | **G** | **110** | **2496** | **G** | **C** | **G** |
| **23** | **939** | **G** | **G** | **A** | **67** | **1861** | **G** | **T** | **T** | **111** | **2513** | **G** | **G** | **C** |
| **24** | **945** | **C** | **C** | **T** | **68** | **1870** | **C** | **G** | **C** | **112** | **2521** | **G** | **A** | **A** |
| **25** | **975** | **C** | **T** | **T** | **69** | **1871** | **T** | **C** | **T** | **113** | **2522** | **A** | **G** | **G** |
| **26** | **1014** | **C** | **T** | **C** | **70** | **1873** | **G** | **T** | **G** | **114** | **2528** | **G** | **G** | **A** |
| **27** | **1038** | **C** | **C** | **G** | **71** | **1874** | **G** | **C** | **G** | **115** | **2533** | **A** | **G** | **A** |
| **28** | **1071** | **C** | **G** | **G** | **72** | **1875** | **T** | **C** | **T** | **116** | **2553** | **G** | **A** | **G** |
| **29** | **1104** | **C** | **T** | **T** | **73** | **1876** | **G** | **C** | **G** | **117** | **2555** | **C** | **T** | **C** |
| **30** | **1135** | **C** | **C** | **A** | **74** | **1878** | **T** | **A** | **T** | **118** | **2564** | **C** | **C** | **G** |
| **31** | **1167** | **G** | **G** | **A** | **75** | **1881** | **C** | **T** | **C** | **119** | **2568** | **A** | **A** | **C** |
| **32** | **1188** | **G** | **C** | **C** | **76** | **1884** | **C** | **T** | **C** | **120** | **2570** | **A** | **A** | **G** |
| **33** | **1202** | **C** | **G** | **G** | **77** | **1886** | **C** | **C** | **G** | **121** | **2573** | **A** | **T** | **G** |
| **34** | **1246** | **C** | **C** | **T** | **78** | **1887** | **T** | **C** | **T** | **122** | **2582** | **T** | **C** | **T** |
| **35** | **1273** | **C** | **C** | **G** | **79** | **1888** | **C** | **G** | **C** | **123** | **2583** | **T** | **C** | **T** |
| **36** | **1276** | **G** | **C** | **C** | **80** | **1889** | **T** | **G** | **T** | **124** | **2591** | **A** | **C** | **A** |
| **37** | **1288** | **G** | **G** | **C** | **81** | **1891** | **G** | **C** | **G** | **125** | **2615** | **C** | **C** | **G** |
| **38** | **1294** | **G** | **C** | **C** | **82** | **1892** | **A** | **T** | **A** | **126** | **2636** | **C** | **T** | **C** |
| **39** | **1300** | **C** | **G** | **C** | **83** | **1900** | **T** | **C** | **T** | **127** | **2657** | **G** | **C** | **G** |
| **40** | **1381** | **C** | **G** | **C** | **84** | **1903** | **G** | **C** | **G** | **128** | **2663** | **A** | **G** | **G** |
| **41** | **1393** | **C** | **C** | **G** | **85** | **1912** | **C** | **C** | **T** | **SNPs** | | | | **128** |
| **42** | **1402** | **C** | **C** | **G** | **86** | **1930** | **C** | **A** | **C** | **Transition** | | | | **47** |
| **43** | **1426** | **A** | **C** | **C** | **87** | **1936** | **G** | **T** | **G** | **Transversion** | | | | **81** |
| **44** | **1438** | **C** | **G** | **G** | **88** | **1955** | **G** | **A** | **A** | **AA Change** | | | | **32** |
