## Supplementary material for "Characterization and Expression Analysis of *Nitrate Reductase 6-1ABD* Gene in Hexaploid Bread Wheat Under Different Nitrogen Regime": Suppl.zip: Table S3.docx

**Table S3: SNPs present between HD 2967 and Choti Lerma for each homoeologues, in *NR 6-1A*, *NR 6-1B,* and *NR 6-1D***

| **Sl. No.** | **Position** | ***HD_NR 6-1A*** | ***CL_NR 6-1A*** | **Sl. No.** | **Position** | ***HD_NR 6-1B*** | ***CL_NR 61-B*** |
| --- | --- | --- | --- | --- | --- | --- | --- |
| **1** | **477** | **T** | **G** | **1** | **12** | **C** | **T** |
| **2** | **1193** | **A** | **G** | **2** | **103** | **G** | **T** |
| **3** | **1199** | **A** | **C** | **3** | **110** | **G** | **C** |
| **4** | **1997** | **A** | **C** | **4** | **120** | **G** | **T** |
| **5** | **2217** | **C** | **A** | **5** | **126** | **C** | **A** |
| **SNPs** | | | **5** | **6** | **127** | **G** | **T** |
| **Transition** | | | **1** | **7** | **133** | **G** | **T** |
| **Transversion** | | | **4** | **8** | **142** | **G** | **A** |
| **AA Change** | | | **3** | **9** | **144** | **C** | **G** |
|  |  |  |  | **10** | **147** | **G** | **A** |
| **Sl. No.** | **Position** | ***HD_NR 6-1D*** | ***CL_NR 6-1D*** | **11** | **151** | **G** | **T** |
| **1** | **750** | **G** | **A** | **12** | **157** | **C** | **A** |
| **2** | **773** | **G** | **A** | **13** | **212** | **A** | **G** |
| **3** | **1173** | **G** | **C** | **14** | **223** | **G** | **A** |
| **4** | **1204** | **G** | **A** | **15** | **226** | **G** | **A** |
| **5** | **1205** | **C** | **A** | **16** | **228** | **G** | **C** |
| **6** | **1206** | **A** | **G** | **17** | **273** | **C** | **G** |
| **7** | **1396** | **G** | **A** | **18** | **278** | **C** | **G** |
| **8** | **2048** | **T** | **C** | **19** | **303** | **G** | **T** |
| **SNPs** | | | **8** | **20** | **369** | **C** | **G** |
| **Transition** | | | **5** | **21** | **389** | **C** | **A** |
| **Transversion** | | | **3** | **22** | **390** | **C** | **T** |
| **AA Change** | | | **5** | **23** | **477** | **G** | **C** |
|  |  |  |  | **24** | **838** | **T** | **A** |
|  |  |  |  | **25** | **480** | **A** | **G** |
|  |  |  |  | **26** | **1181** | **C** | **A** |
|  |  |  |  | **27** | **1464** | **A** | **G** |
|  |  |  |  | **28** | **2707** | **C** | **G** |
|  |  |  |  | **SNPs** | | | **28** |
|  |  |  |  | **Transition** | | | **7** |
|  |  |  |  | **Transversion** | | | **21** |
|  |  |  |  | **AA Change** | | | **24** |
