## Supplementary material for "Characterization and Expression Analysis of *Nitrate Reductase 6-1ABD* Gene in Hexaploid Bread Wheat Under Different Nitrogen Regime": Suppl.zip: Table S4.docx

**Table S4: Changes in amino acid in NR 6-1 protein due to SNPs among different homeologues of wheat and also between HD 2967 and Choti Lerma for each homoeologues**

| **AA Changes among the homeologues** | | | | | **AA Changes between HD and CL** | | | |
| --- | --- | --- | --- | --- | --- | --- | --- | --- |
| **Sl.No.** | **Position** | ***NR 6-1A*** | ***NR 6-1B*** | ***NR 6-1D*** | **S.l.No.** | **Position** | ***HD_NR 6-1A*** | ***CL_NR 6-1A*** |
| 1 | 32 | S | G | S | 1 | 159 | F | V |
| 2 | 33 | A | G | G | 2 | 665 | E | D |
| 3 | 130 | R | K | K | 3 | 739 | L | I |
| 4 | 149 | K | K | R | **SI.No.** | **Position** | ***HD_NR 6-1B*** | ***CL_NR 6-1B*** |
| 5 | 194 | I | I | V | 1 | 1 | M | C |
| 6 | 427 | M | M | I | 2 | 34 | G | C |
| 7 | 515 | A | T | T | 3 | 36 | R | T |
| 8 | 520 | F | Y | Y | 4 | 42 | D | Y |
| 9 | 596 | T | A | T | 5 | 44 | D | Y |
| 10 | 605 | S | N | S | 6 | 47 | D | K |
| 11 | 606 | G | A | A | 7 | 50 | G | C |
| 12 | 616 | P | S | A | 8 | 52 | P | T |
| 13 | 617 | A | V | A | 9 | 70 | K | R |
| 14 | 618 | G | V | G | 10 | 74 | D | N |
| 15 | 623 | V | P | V | 11 | 75 | E | N |
| 16 | 624 | L | Q | L | 12 | 92 | T | S |
| 17 | 625 | P | L | P | 13 | 100 | E | D |
| 18 | 626 | P | L | P | 14 | 122 | N | K |
| 19 | 627 | L | P | V | 15 | 129 | A | D |
| 20 | 628 | S | A | S | 16 | 158 | Q | H |
| 21 | 629 | T | S | T | 17 | 279 | F | L |
| 22 | 650 | V | M | M | 18 | 393 | T | D |
| 23 | 666 | H | Q | Q | 19 | 402 | C | E |
| 24 | 698 | D | E | E | 20 | 735 | D | E |
| 25 | 704 | E | E | D | 21 | 736 | I | S |
| 26 | 730 | A | S | S | 22 | 737 | K | R |
| 27 | 760 | P | R | R | 23 | 738 | G | A |
| 28 | 800 | Y | Y | F | 24 | 740 | V | W |
| 29 | 830 | V | L | V | **S.l.No.** | **Position** | ***HD_NR 6-1D*** | ***CL_NR 6-1D*** |
| 30 | 838 | G | E | E | 1 | 250 | D | N |
| 31 | 848 | D | N | D | 2 | 391 | V | L |
| 32 | 855 | D | E | D | 3 | 401 | G | E |
|  |  |  |  |  | 4 | 402 | L | V |
|  |  |  |  |  | 5 | 465 | G | D |
