## Supplementary figures and images for "Characterization and Expression Analysis of *Nitrate Reductase 6-1ABD* Gene in Hexaploid Bread Wheat Under Different Nitrogen Regime"

### Fig. S2.jpg

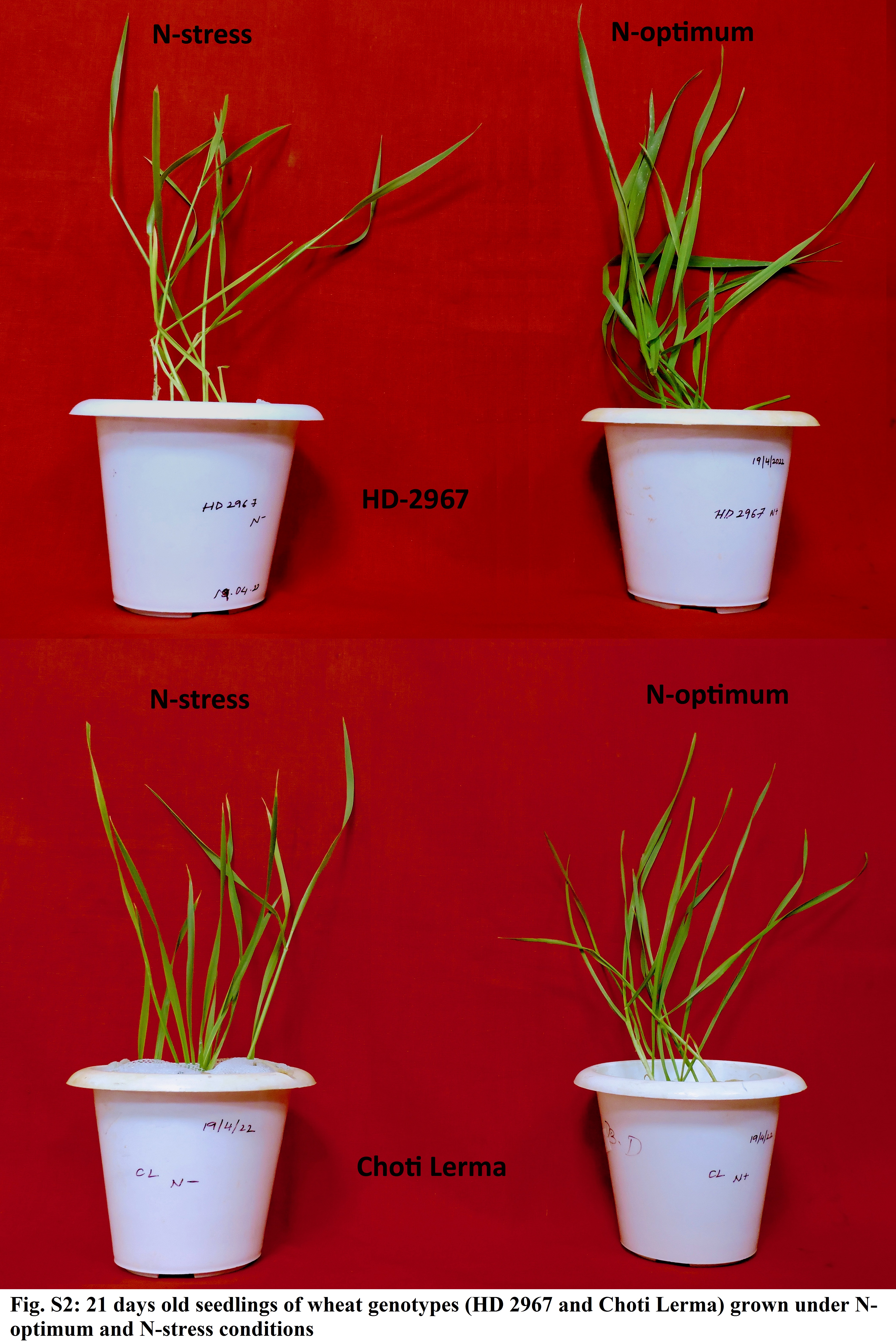

### Fig. S3 MSA of NR 6-1 CLONED CDS.jpg

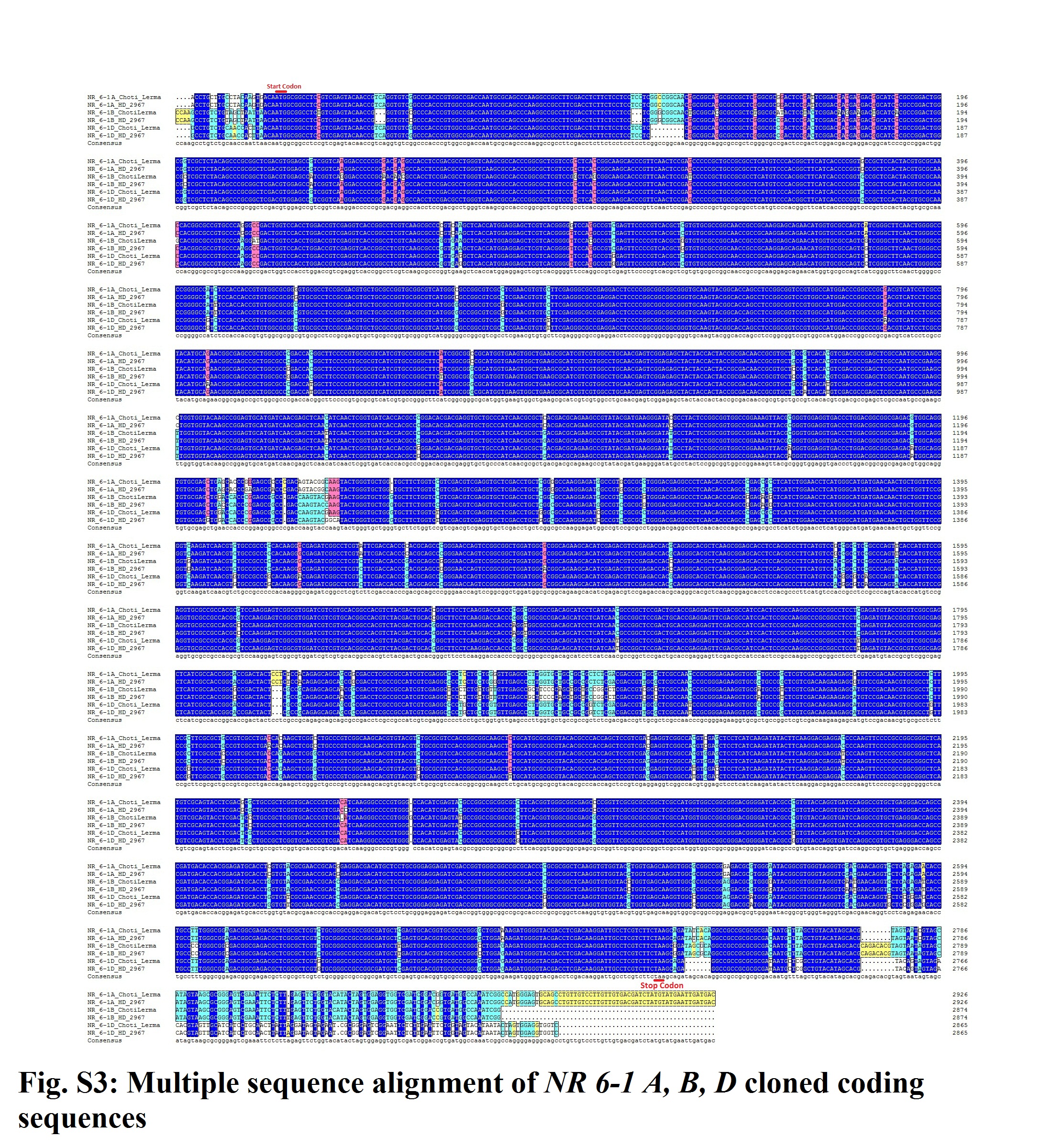

### Fig.S1 cis-elements.jpg

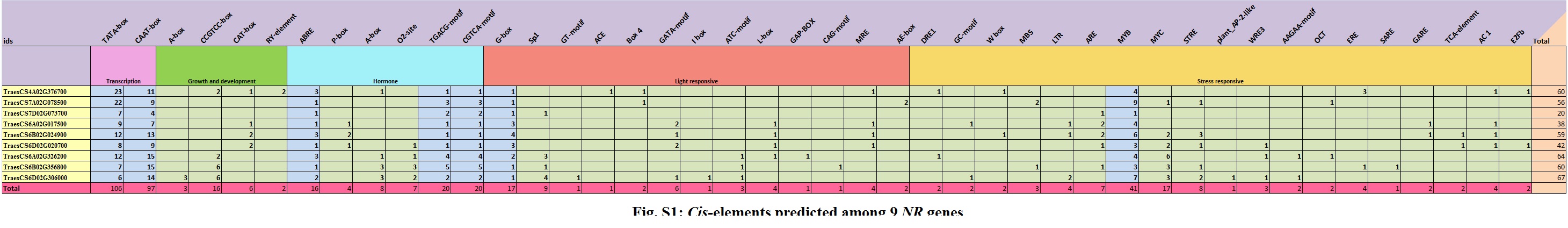
